# Boosting of neural circuit chaos at the onset of collective oscillations

**DOI:** 10.1101/2022.08.28.505598

**Authors:** Agostina Palmigiano, Rainer Engelken, Fred Wolf

## Abstract

Neuronal spiking activity in cortical circuits is often temporally structured by collective rhythms. Rhythmic activity has been hypothesized to regulate temporal coding and to mediate the flexible routing of information flow across the cortex. Spiking neuronal circuits, however, are non-linear systems that, through chaotic dynamics, can amplify insignificant microscopic fluctuations into network-scale response variability. In nonlinear systems in general, rhythmic oscillatory drive can induce chaotic behavior or boost the intensity of chaos. Thus, neuronal oscillations could rather disrupt than facilitate cortical coding functions by flooding the finite population bandwidth with chaotically-boosted noise. Here we tackle a fundamental mathematical challenge to characterize the dynamics on the attractor of effectively delayed network models. We find that delays introduce a transition to collective oscillations, below which ergodic theory measures have a stereotypical dependence on the delay so far only described in scalar systems and low-dimensional maps. We demonstrate that the emergence of internally generated oscillations induces a complete dynamical reconfiguration, by increasing the dimensionality of the chaotic attractor, the speed at which nearby trajectories separate from one another, and the rate at which the network produces entropy. We find that periodic input drive leads to a dramatic increase of chaotic measures at a the resonance frequency of the recurrent network. However, transient oscillatory input only has a moderate role on the collective dynamics. Our results suggest that simple temporal dynamics of the mean activity can have a profound effect on the structure of the spiking patterns and therefore on the information processing capability of neuronal networks.

---

Cortical spiking patterns are unreliable and vary from trial to trial in response to identical stimuli (1, 2). Multiple factors contribute to this noise entropy – variability unrelated to the stimulus or to the network state – from network-based deterministic chaos (3–6) to stochasticity in the action potential generation (7) and transmission (8). Noise entropy necessarily interferes with the robust representation of spiking patterns and code-word based information flow, and poses the question of which possible mechanisms cortical networks implement to represent and transmit information in the presence of this undesired variability. One possibility is that cortex integrates each-cell spikes in a few ms window to represent stimuli. In this rate code, spurious changes in spiking patters contribute to asynchronous irregular dynamics with low correlations. However, if precisely-timed spike patterns are the code used by cortical circuits, spiking variability is an undesired feature, and other network mechanisms should exist to control it.

Neuronal oscillations have been proposed to be a backbone organizing neuronal firing (9) that presumably quenches spike-pattern variability by reducing the number of possible network configurations. Transient oscillatory (but not strongly synchronous) activity can aid the flexible gating information flow (10), but whether any type of oscillatory activity is able to tame noise entropy is still an unanswered question. In fact, from a theoretical perspective, whether noise entropy due to chaotic dynamics is intensified or quenched by collective oscillations is not easily predictable. First, the internally-generated population oscillation can be thought of as a common harmonic drive to all the cells in the network. Generally, whether a harmonic drive will intensify or quench chaos in a given system is not possible to know a priory, and both effects have been observed (11–14). Second, oscillatory activity has been theoretically (15–20) and experimentally (21–24) traced back to delayed recurrent inhibition, whose effect is also difficult to predict: On the one hand, delayed interactions can induce highly synchronized and therefore highly structured activity (25–27), possibly shrinking the dimensionality of the dynamics and quenching variability. On the other hand, the dimensionality of the chaotic attractor increases linearly with increasing delays (28, 29) in simple systems, and it is possible that similar dependencies occur in more complex systems than the ones studied so far.

Here we investigate whether the emergence of collective oscillations at the population level quenches noise entropy by synchronizing neuronal firing or contributes to an expansion of the number of the possible spike-pattern configurations explored by the circuit by intensifying chaos. To do so, we advance the tractability of large neuronal spiking networks of exactly solvable neuronal models by developing a strategy that allows analyzing the dynamics of delayed spiking networks in a system with fixed and finite degrees of freedom. By building a two-stage model of the action potential generation and transmission, we map the infinite-dimensional system to a finite one and semi-analytically compute the entire spectrum of Lyapunov exponents whose positive sum is a direct proxy for the network’s noise entropy (6). We find that when delays are small, the dimensionality of the attractor increases linearly with the delay while keeping noise entropy constant, analogously to what has been reported in simplified delayed systems (28, 29). Larger delays destabilize the asynchronous irregular dynamics of the spiking network, giving rise to population oscillations in the beta frequency range (∼ 20 Hz). We show that this transition is achieved via Hopf bifurcation of the mean activity at a critical set of parameters that can be computed exactly, and is analogous to what is reported in network models without a dynamical mechanism for action potential generation (15–18). Unexpectedly, the noise entropy, together with the dimensionality of the chaotic attractor, is boosted at the oscillatory instability. To investigate the mechanism that underlies this effect, we drive a network with a periodic external input, to first order, analogous to the one that the network receives at the oscillatory transition. We find that the external periodic drive dramatically boosts noise entropy in the neighborhood of the resonant frequency of the network, and quenches it otherwise. We find that a realistic drive, in which collective activity has short transients of oscillatory activity with drifting frequencies, leads to only moderate changes in the noise entropy. Our results indicate that simple dynamics of the mean activity can have a profound effect on the chaotic dynamics of the network and that realistic transient synchrony signals could, beyond flexibly gating information transmission, prevent boosting of undesired variability in the cortex.

## Results

### Delayed networks with a finite-dimensional phase space

Spiking networks have an innate source of instability, the generation of the action potential, which leads to chaotic dynamics even when the statistics of the population firing rate are independent of time (3, 6). The noise entropy generated by these networks, present even in the absence of any stochastic source of variability, can be computed through the spectrum of Lyapunov exponents (6) (see *Methods*). Therefore, given a group of control parameters that control the transition to oscillations in the network we can ask: does the onset of collective rhythms imply a reduction of the intensity of chaos and a more robust representation of the network’s inputs? Or does it imply the opposite?

One well-known parameter controlling the transition to collective oscillations in brain-like inhibitory-dominated networks is the synaptic delay (30, 31) (Fig. 1 **a-b**). This parameter, despite being inescapable in biology, introduces a new degree of complexity: even simple scalar differential equations (Fig. 1**c**) are transformed into an infinite-dimensional system when incorporating delayed feedback (Fig. 1**d**). Whether a discrete spectrum of Lyapunov exponents generally exists in delayed systems, and whether it can be computed exactly, remains poorly understood. The most general strategy proposed so far suggests binning the delay period and expanding the dimensionality of the system by the number of bins considered (32). However, this procedure becomes arbitrarily costly with increasing resolution, even in scalar equations.

**Fig. 1.**
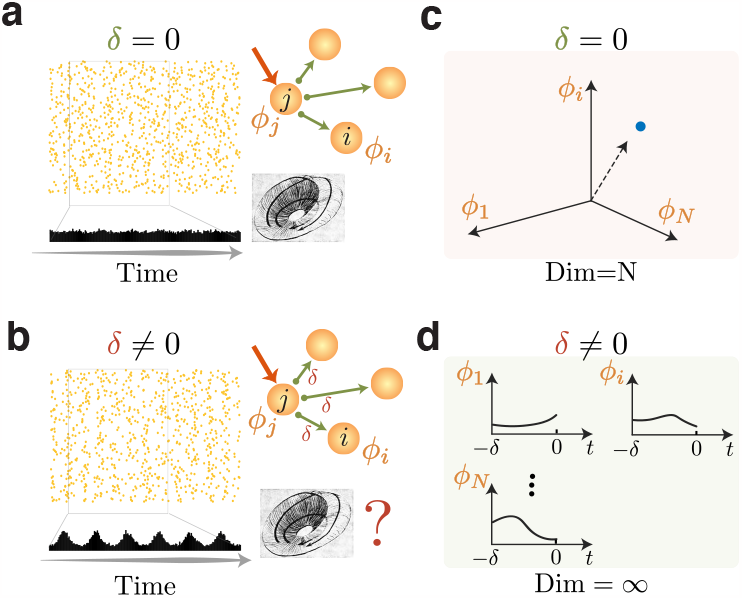
Fundamental challenges of the study of oscillations due to delayed interactions. **(a)** Neuronal network without synaptic delays. Whenever the unit with dynamical variable *ϕ*_*j*_ spikes, its spike is immediately relayed. **(b)**. Delayed network. Each time a neuron *j* is active, it takes a time *δ* for the spike to propagate to the postsynaptic neurons. In the case of delayed pulse-coupled oscillators, this network can be mapped to a finite size system, with variable dimension (33). **(c)** An *N* dimensional deterministic dynamical system is defined by *N* scalar initial conditions.**(d)** In a delayed dynamical system, each of the *N* variables will be initialized by a history function. In this case, the system will be infinite-dimensional.

We developed a strategy that allows for studying the impact of collective oscillations on the properties of the network’s chaos using delayed recurrent inhibition as the control parameter in a numerically exact and efficient way. We include delays dynamically by designing a two-stage single-neuron model composed of a soma, modeled as a quadratic integrate-and-fire neuron (*ϕ*), and a single compartment axon (SCA, *ξ*, see Fig. 2**a**, *Methods* and Fig. S1 of the *Supplementary Information*). The SCA is modeled as an excitable spiking unit, resting in a steady state, which depolarizes at spike arrival. Post-synaptic delays are introduced as the time it takes for the axon to spike and relay its input. This strategy allows mapping the infinite-dimensional system to a system with 2*N* dimensions, under the assumption that the interspike interval is much larger than the delay *δ*. Numericallyexact event-based simulations of a network of inhibitory neurons with short delays (see below for E-I networks) exhibit the characteristic dynamics of non-delayed balanced state networks, with asynchronous irregular spiking patterns (Fig. 2**b**), broad firing rate distributions (Fig. 2**c**) and a large coefficient of variation of the inter spike intervals (Fig. 2**d**).

**Fig. 2.**
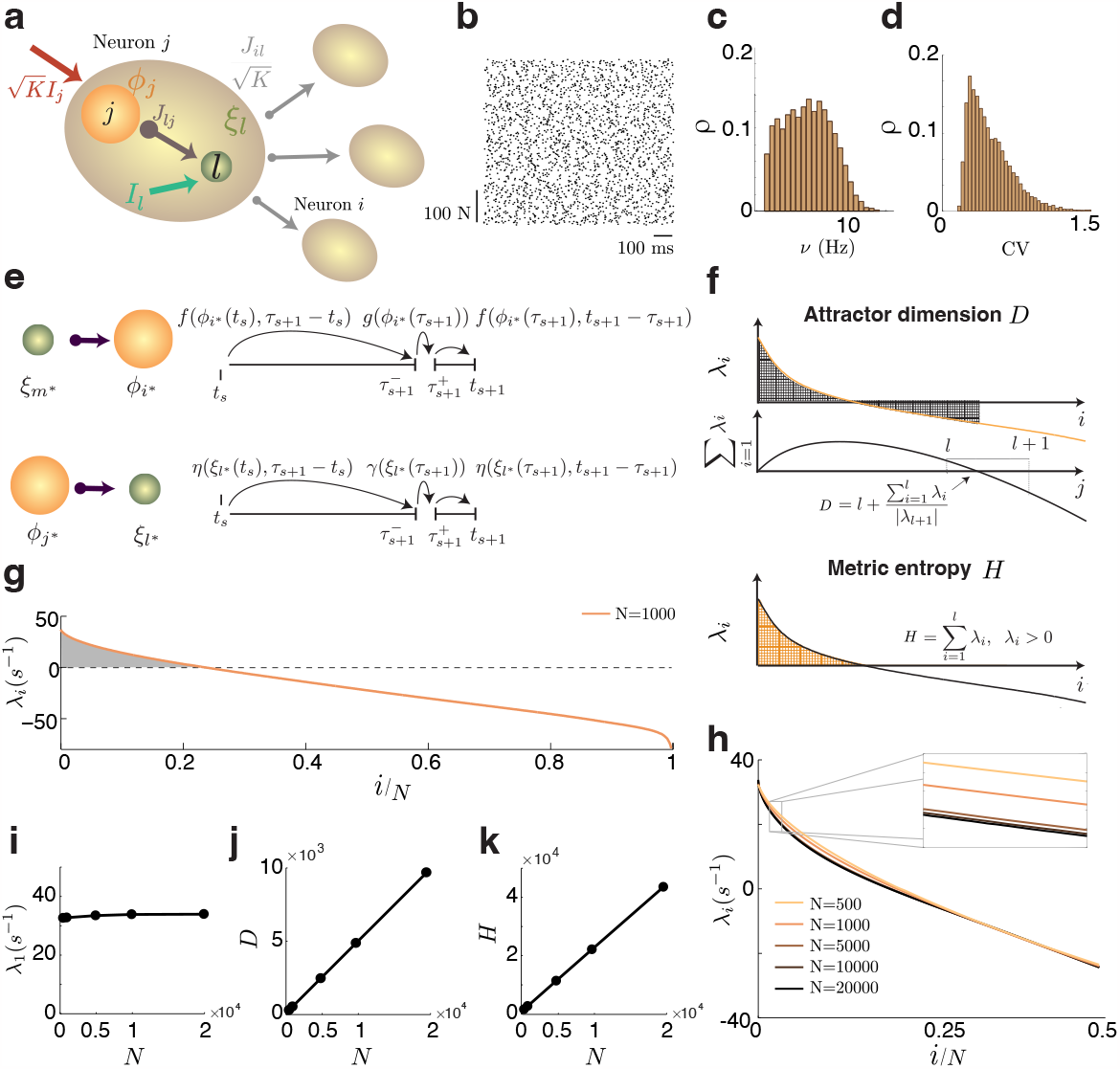
Extensive chaos in delayed balanced circuits. **(a)** Circuit diagram of the two-stage neuronal model. Each neuron (brown) is composed of a soma (orange) linked to its single-compartment-axon (SCA, green). The link weight *J*_*lj*_ is such that when the soma spikes (i.e. *ϕ*_*j*_ reaches threshold), the SCA (*ξ*_*l*_) is released from its fixed-point steadystate, relaying its input after a delay *δ* exactly defined by their connection weight (see Eq. 10 *Methods*). Each soma receives on average the activity of *K* neighbors with weight 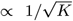, and external inputs of order 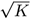. **(b)** Raster plot of network’s activity. **(c)** Firing rate distribution (left) and **(d)** CV distribution (right) for a small (1 ms) delay. **(e)** Terms involved in the calculation of the single-spike Jacobian. When an SCA (soma) spikes, the state of the postsynaptic soma (SCA) is propagated via the function *f* (*η*) from the previous spike time *t*_*s*_ to the new spike time *τ*_*s*+1_, then updated via *g* (*γ*), and then propagated up to the next spike time. Then the limit of *τ*_*s*+1_ → *t*_*s*+1_ is taken (see *Methods*). **(f)** Chaos descriptors: Kaplan-York dimension (top) and the metric entropy (bottom). **(g)** Spectrum of Lyapunov exponents. Shaded area are the positive exponents. **(h)** Spectrum for different network sizes as a function of the normalized index of the exponent. A system-size invariant Lyapunov spectrum indicates extensive chaos **(i)** First Lyapunov exponent **(j)** Attractor dimension **(k)** and Metric entropy as a function of system size (N).

We compute the entire spectrum of Lyapunov exponents by deriving an analytical expression of the Jacobian of the effectively delayed network that is evaluated numerically at each spike time. The Jacobian’s non-trivial elements will characterize either the change in the state of a post-synaptic soma to an infinitesimal change in the state of the pre-synaptic SCA (Fig. 2**e** top) or the change in the state of an SCA to an infinitesimal change in the state of the pre-synaptic soma (Fig. 2**e** bottom, see *Methods* and *Supplementary Information* for an in-depth description).

The spectrum of exponents is shown in Figure 2**g** (see Fig. S2 of the *Supplementary Information* for different parameter choices of the SCA and S3 for convergence). The spectrum is invariant of network size (N), hinting at a continuous density of Lyapunov exponents for infinitely large networks (Fig. 2**h**), and a network-size independent first Lyapunov exponent (Fig. 2**i**). The attractor dimension (Fig. 2**j**) and the metric entropy, a proxy for the noise entropy in the network (6) (Fig. 2**j**), grow linearly with network size corresponding to a network-size invariant spectrum, which invites defining *intensive* measures of dimension 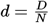 and metric entropy 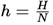, the latter in units of bits per second.

### Boosting and collapse of chaos

As expected, sufficiently long delays destabilize the asynchronous irregular state in balanced-state networks of QIF neurons and lead to a state of collective oscillations in which the irregular dynamics at the single cell level is preserved (Fig. 3**a**). The mean square deviation (MSD, see *Methods*) of the mean firing rate has a threshold-linear dependence on the delay, which is the signature of a Hopf bifurcation at the level of the population mean, and is consistent with previously described oscillatory transitions to analogous states in related systems (15, 18, 19, 31, 34). The critical delay *δ*_*MSD*_ that indicates the onset of oscillatory activity depends on the parameters of the network, decreasing with increasing connection strength and target mean firing rate, and becoming arbitrarily small for large in-degree (see below). A second instability, now to full network synchrony, occurs at a second critical delay. The collective oscillation, in the beta range up to that point, collapses to a fully synchronous state with a frequency defined by the network’s target mean firing rate of 5 Hz (Fig. 3**c**).

**Fig. 3.**
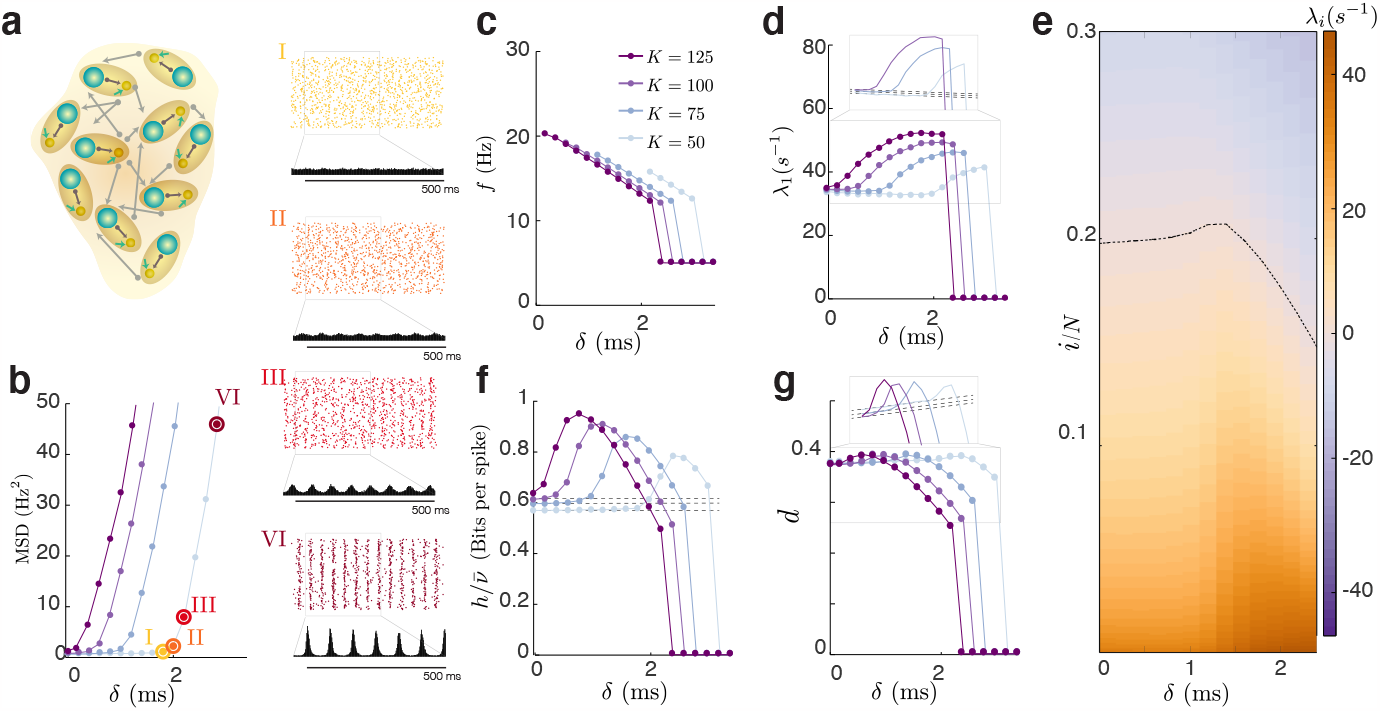
Boosting and collapse of chaos with increasing delayed inhibition. **(a)** Diagram of an inhibitory network with dynamic delays, each neuron is as in Fig. 2a. The mean firing rate of the network is kept constant at 5 Hz by adjusting the homogeneous external input that each neuron receives. **(b)** Left: Mean squared deviation from the mean population firing rate (MSD, vanishing values indicates an asynchronous state) as a function of the synaptic delay for different values of the average in-degree *K*. The larger *K*, the smaller the delay *δ* needed to destabilize the asynchronous irregular state. Numbered circles indicate the points for which raster plots are shown. **(c)** Frequency of the oscillation is in the beta range and monotonically decreases with the delay. After a *K*-dependent critical delay *δ*_*c*_, the network collapses to a limit cycle (complete synchrony), and the network oscillates at the target firing rate *ν* (see *Methods*) **(d)** Maximum Lyapunov exponent as a function of *δ* for different values of *K*. **(e)** A third of the spectrum of Lyapunov exponents as a function of the delay for *K* = 75. The blue line in panel (d) is the bottom row of this panel. **(f)** Intensive metric entropy (*h* = *H/N*) divided by the mean firing rate 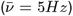, giving units of bits per spike, and **(g)** attractor dimension *d* = *D/N*, as a function of the delay. Both the metric entropy and the attractor dimension have a non-monotonic dependence with the delay prior to the collapse to a limit-cycle.

Surprisingly, we find that contrary to the intuition of increased regularity in oscillatory networks, chaos is boosted at the (first) oscillatory transition. The maximum Lyapunov exponent (LE) decreases monotonically with increasing delays up to the oscillatory instability (Fig. 3**d**) together with around 6% of the LEs (Fig. 3**e**, see also Fig. S4 of the *Supplementary Information*). Simultaneously, the number of positive LEs increases monotonically (Fig. S4**a**). These competing effects contribute to the remarkable fact that the metric entropy remains independent of the delay up to until the oscillatory instability (Fig. 3**f**, compare to dashed line). The attractor dimension, on the other hand, is dominated by the number of positive LEs and increases monotonically up to the transition (Fig. 3**g**). Interestingly, up to the oscillatory instability the ergodic properties of the pulse-coupled highdimensional network studied here match the dependencies reported for low-dimensional systems (and also the dependences of rate networks, see below); with increasing delay the maximum Lyapunov exponent decreases, the attractor dimension increases, while the metric entropy is independent of the delay (28, 35, 36).

At the critical delay, all chaotic measures intensify. Looking at a fraction of the Lyapunov spectrum as a function of the delay (Fig. 3**e**) reveals that i) all the positive LEs increase in value after the transition and ii) the total number of positive LEs initially increases monotonically after the first oscillatory transition but later decreases (Fig. S4**a**). The entropy and the attractor dimension at the transition can only increase given the increase in the magnitude and the amount of positive LEs (Fig. 3**f**,**g**). Although the positive LEs grow larger in magnitude, their number decreases soon after the transition; the uneven contribution of these opposing effects gives rise to the non-monotonic behavior of the entropy and the attractor dimension. These dependencies are not a finite size effect (see Fig. S3**d**) and we have not observed signs of bistability (37).

We compare the original, infinite-dimensional delayed system with the equivalent finite system studied here by self-consistently computing the instability periphery in parameter space (following Ref. (38), see *Methods*). We compute the critical value of the delay at which the mean firing rate (in the original, delayed system) undergoes a Hopf bifurcation, and we compare it to the one obtained numerically by estimating the critical delay from the MSD’s dependency on the delay, *δ*_*MSD*_ (Fig. 4**c**) the first LE, *δ*_*LE*_, (Fig. 4**d**) and the metric entropy, *δ*_*H*_, (Fig. 4**e**). Concretely, we compute the delay value at which the derivative of the maximum LE with respect to the delay changes from negative to positive, the delay at which the derivative of the metric entropy departs from zero, and the delay at which the derivative of the attractor dimension departs from a constant value. The accuracy of the match in panels (*c-e*) depends on both the estimation of the critical parameters from ergodic theory quantities, which are sensitive to simulation size (see Fig. 4), and details of the implementation of the self-consistent estimation of the critical delay in the original system (see also Fig. S5 in the *Supplementary Information*). Despite the assumption of fixed in-degree implicit in the semi-analytic method used here to compute the critical delay (38), broken in our simulations, we find good agreement.

**Fig. 4.**
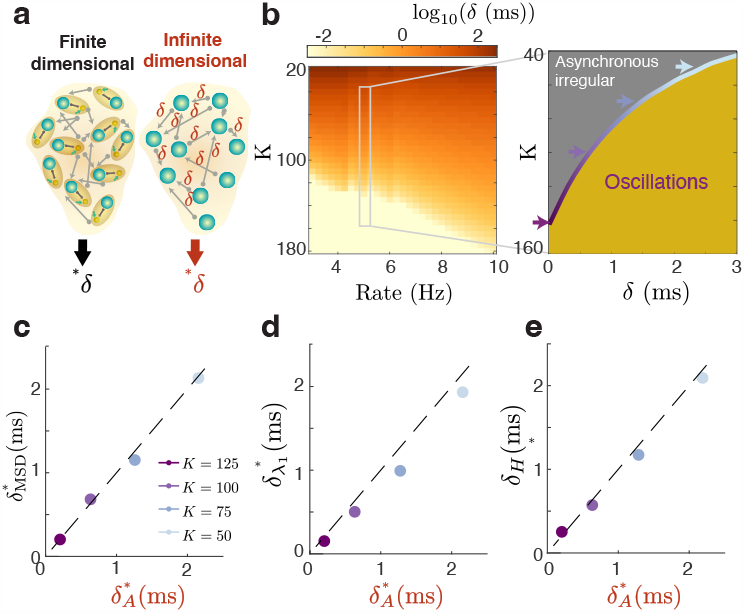
Chaos is boosted at the oscillatory instability. **(a)** We compare the infinite-dimensional system (original delayed system) with the system with fixed and finite degrees of freedom studied here. **(b)** Phase diagram (computed semi-analytically following (38)) showing the critical delay at which the asynchronous irregular state loses stability to collective oscillations, as a function of the mean degree K and the target mean firing rate. **(c)** Comparison between the critical delay estimated from simulations as when the MSD departs from zero (*δ*_*MSD*_) and the one from the true delayed system (*δ*_*A*_) **(d)** Same as c) but for the critical delay estimated in simulations is estimated from when the first Lyapunov exponent increases 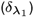 **(e)** Same as c) but for the critical delay estimated from the increase in the metric entropy (*δ*_*H*_)

Does the emergence of collective oscillations always lead to boosting of chaos? We test two other mechanisms that induce oscillations in the network. First, as suggested in Fig. 4, increasing the average degree (K) can induce oscillations in this network even in the absence of delayed interactions. *Supplementary Figure* S6 demonstrates that all indicators of chaos (*λ*_1_, *d* and *h*) also increase at the oscillatory transition as a function of the mean degree K. Second, we find that in non-delayed excitatory-inhibitory networks, there is a transition to a balanced oscillatory regime with an increasing inhibitory time constant, in agreement to what is reported in binary networks (39). We find that, right after the transition to collective oscillations, all indicators of chaos are intensified. Taken together, our results suggest that the transition from an asynchronous irregular activity to collective oscillations, although only slightly changing the single neuron spiking statistics, induces a full reconfiguration of the network dynamics by simultaneously increasing the maximum Lyapunov exponent, the metric entropy and the dimension of the chaotic attractor.

### Single-cell heterogeneity perpetuates chaos

In a network in which all the neurons are the same and receive the same inputs, the only source of disorder is from the sparse connectivity. Increasing delays in these types of networks pushes neurons to align their activity to the ongoing rhythm until the dynamics of the network collapses to a limit cycle (Fig 3, *δ* ≈ 2.1 ms).

How robust is this pathological state of full synchrony? Would it be observable in cortex where cell and cell-type heterogeneity are abundant? We find that the inclusion of even moderate amounts of single-cell heterogeneity not only curbs the synchronization capability of the network (16, 40, 41) (Fig. 5**a**), but also safeguards a biophysically plausible regime by eliminating the second oscillatory instability leading to a pathological full-synchrony state (Fig.5**b**). In this case, the maximum LE (and also the coefficient of variation, see Fig. S9 and S10 of the *Supplementary Information*) peaks at the delay that would have led to a full-synchrony state in the absence of heterogeneity. The network, instead of collapsing to a limit cycle, retains the irregular firing patterns and the microscopic chaos associated with it for delays as long as the membrane time constant (Fig.5**b-h**, see also Fig. S8 in the *Supplementary Information*). The ghost of the second oscillatory transition marks a regime in which increasing delays can only further align the neurons to the ongoing rhythm, resulting in a vanishingly small metric entropy for large delays (Fig.5**f**). Interestingly, for large delays the maximum LE can have the same value as with no delay while the metric entropy undergoes a ten-fold decrease, indicating that simple numerical estimates of the maximum LE could not reveal the full extent of the dynamical reconfiguration the network undergoes after the second oscillatory transition. These results are robust at all network parameters (the connection strength, the target mean firing rate, and the average degree K) as shown in Figures S9 and S10 of the *Supplementary Information*.

**Fig. 5.**
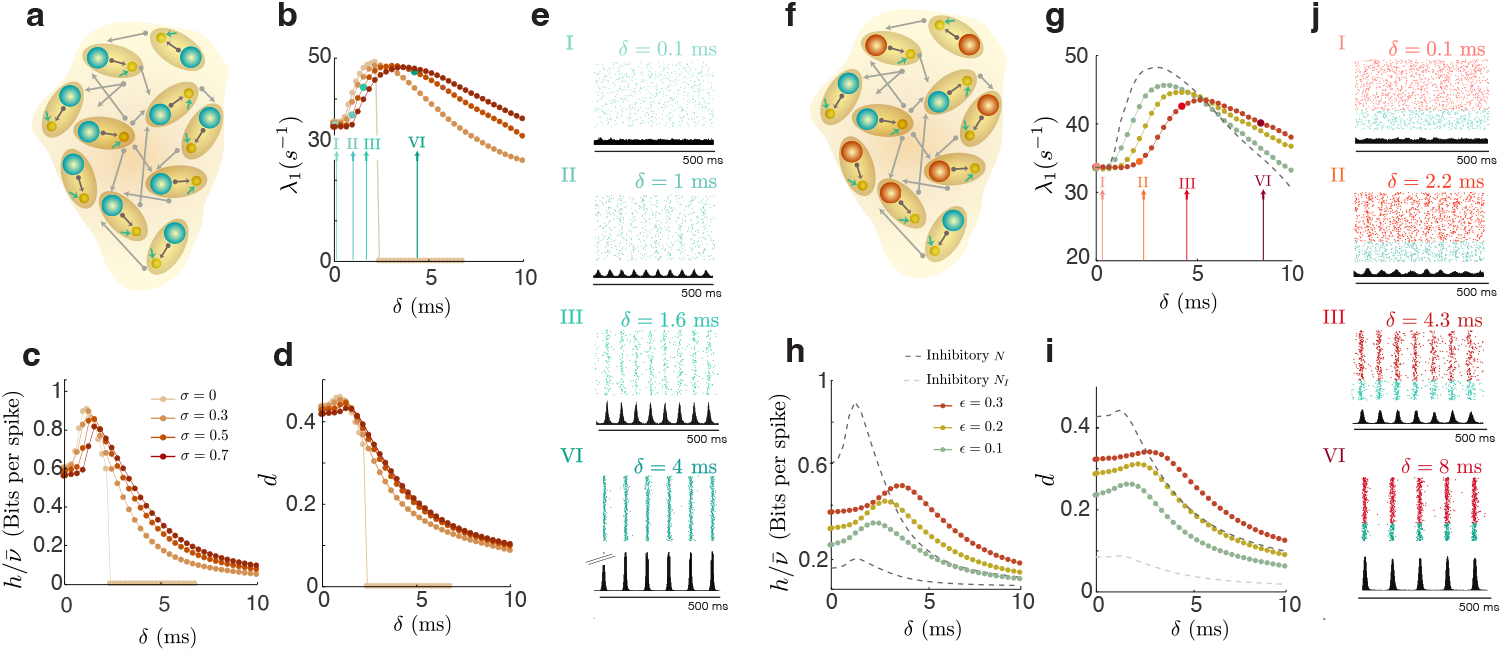
Single cell and cell-type heterogeneity perpetuate chaos. **(a)** Network of heterogeneous inhibitory neurons. **(b)** Maximum Lyapunov exponent **(c)**, network-size intensive metric entropy normalized by the firing rate and **(d)** intensive attractor dimension as a function of the delay for different values of heterogeneity. The homogeneous case (of Fig. 3) is shown for reference. **(e)** Spiking patterns at different values of the synaptic delays. Notice how despite the seemingly strong synchronization for large delays, the network remains strongly chaotic. **(f)** Diagram of a network of *N* neurons, *N*_*E*_ = 0.8*N* excitatory and *N*_*I*_ = 0.2*N* inhibitory, in the balanced state. The connectivity matrix is proportional to 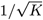 with entries *J*_*EE*_ ∝ *ϵη, J*_*IE*_ ∝ *ϵη*, 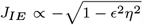, and 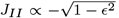, and *η*=0.9 **(g)** Maximum LE as a function of the delay for different strength of the excitatory to inhibition connectivity *ϵ*. Dashed dark gray is a network of *N* inhibitory neurons. **(h)** Intensive metric entropy as a function of the delay for different values of *ϵ* and for a network of purely inhibitory neurons of size *N* (dashed dark gray) and *N*_*I*_ (dashed light gray). **(i)** Intensive attractor dimension as a function of the delay for different values of *ϵ* and for a network of purely inhibitory neurons of size *N* (dashed dark gray) and *N*_*I*_ (dashed light gray). **(j)** Spiking patterns. Spikes in red are of E neurons, while blue are of I.

The presence of strong recurrent excitation introduces a new source of network instability. Stable cortical circuits that would be unstable in the absence of feedback inhibition are called inhibition stabilized (42, 43), with balanced networks being one subclass of those. We next asked whether this more general class of model would also exhibit boosting of chaos at the oscillatory transition. We study the two cell-type case by including recurrent excitation in a four-to-one ratio, as observed in cortex(44) (Fig. 5**e**). We require identical input currents statistics to inhibitory and excitatory neurons (see *Methods*), and identical delays within and across the populations. To guarantee the existence of a balanced state, we used a parametrization (3) that relies on the strength of the E to I connections, *ϵ*. We find that the strength of the EI loop has a stabilizing effect on the asynchronous irregular regime, requiring increasingly longer delays for the oscillatory instability to occur. We notice that when the inputs to the different cell-type populations are identical, like in this case, the network synchronizes without a lag between the E and I populations (19). By replacing 80% of the neurons with excitatory neurons, the network is comparatively less chaotic for moderate delays (Fig. 5**g-i**, compare dashed dark-grey line with colored ones). On the other hand, for a fixed amount of inhibitory neurons, increasing the intensity of the E-I feedback loop *ϵ* intensifies chaos for all delays (Fig. 5 **g-i**, compare dashed light-grey line with colored ones, see also Fig. S11 of the *Supplementary Information*).

### Absence of oscillatory boosting in rate networks

Next, we asked whether rate networks, a fundamentally different type of neuronal network model, would exhibit a transition to collective oscillations with delayed interactions, and if so, whether this transition would lead to an increase in noise entropy. We analyze a variety of both classic (45) and purely inhibitory rate networks (46), which we name weakly-delayed rate networks. We incorporate delays perturbatively by expanding the delayed activity around the non-delayed activity in powers of the delay up to order *k* (𝒪 (*k*)) (see *Methods*, and Section 3 of the *Supplementary Information*). By incorporating this expansion, we increase the degrees of freedom of a non-delayed network by *N* 𝒪 (*k*), where N is the number of neurons. A scheme of the effective network after a second order expansion is shown in *Supplementary Figure* S13**a**.

We analyze the dependence on the delay for the balanced inhibitory network and find that, although it has a transition from a chaotic regime (Fig. S14**a** top panel) to clock-like synchrony (Fig. S14**a** bottom) after a critical delay as in the spiking network without heterogeneity, we did not observe a coexistence of chaos with a collective oscillation at intermediate delays (unlike (47) under the presence of feedback). We computed the entire spectrum of Lyapunov exponents of the classic (Fig. S13) and the inhibitory network (Fig. S14) in the standard way (see Ref. (48)) as a function of the synaptic delay. We find that, before the collapse to a limit cycle in the inhibitory network and for all the delays analyzed in the classic network, the maximum Lyapunov exponent decreases (Figs. S13**e** and S14**c**), the metric entropy is constant (Figs. S13**g** and S14**e**) and the attractor dimension increases linearly with the delay (Figs. S13**f** and S14**d**), as we observed in spiking networks before the first oscillatory instability, and as was described in scalar differential equations and low dimensional maps (28, 29).

### Resonant boosting of chaos

If the mere existence of collective oscillations was a sufficient condition to observe an intensification of all chaotic measures, then driving the network with an inhomogeneous Poisson process, harmonically modulated at frequencies smaller than the cutoff (38, 49), should induce oscillations at the population level and intensify chaos as quantified by all ergodic measures described above.

To test this hypothesis, we first include an external harmonic drive to a non-delayed purely inhibitory network. Each neuron receives, besides the recurrent input, an external input composed of K spike-trains that are sampled from an inhomogeneous Poisson process with a harmonically modulated rate with baseline *I*_0_, amplitude *I*_1_, and frequency *f* (Fig. 6**a**, left). We find that increasing the amplitude of the harmonic drive successfully induces population oscillations (Fig. 6**a**, right panels) as apparent from the raster plots. Interestingly, for low frequencies, even if there is a collective rhythm, chaos is not intensified but is reduced instead. The larger the amplitude of the oscillatory modulation of the incoming spike-trains, the more strongly reduced the first Lyapunov exponent (Fig. 6**b**), the metric entropy (Fig. 6**c**) and the attractor dimension are (Fig. 6**d**). Reduction of chaos by an external noisy input has been previously observed in very similar systems to the one studied here (6) and has been extensively characterized both analytically and numerically in rate networks (14, 48, 50–52).

**Fig. 6.**
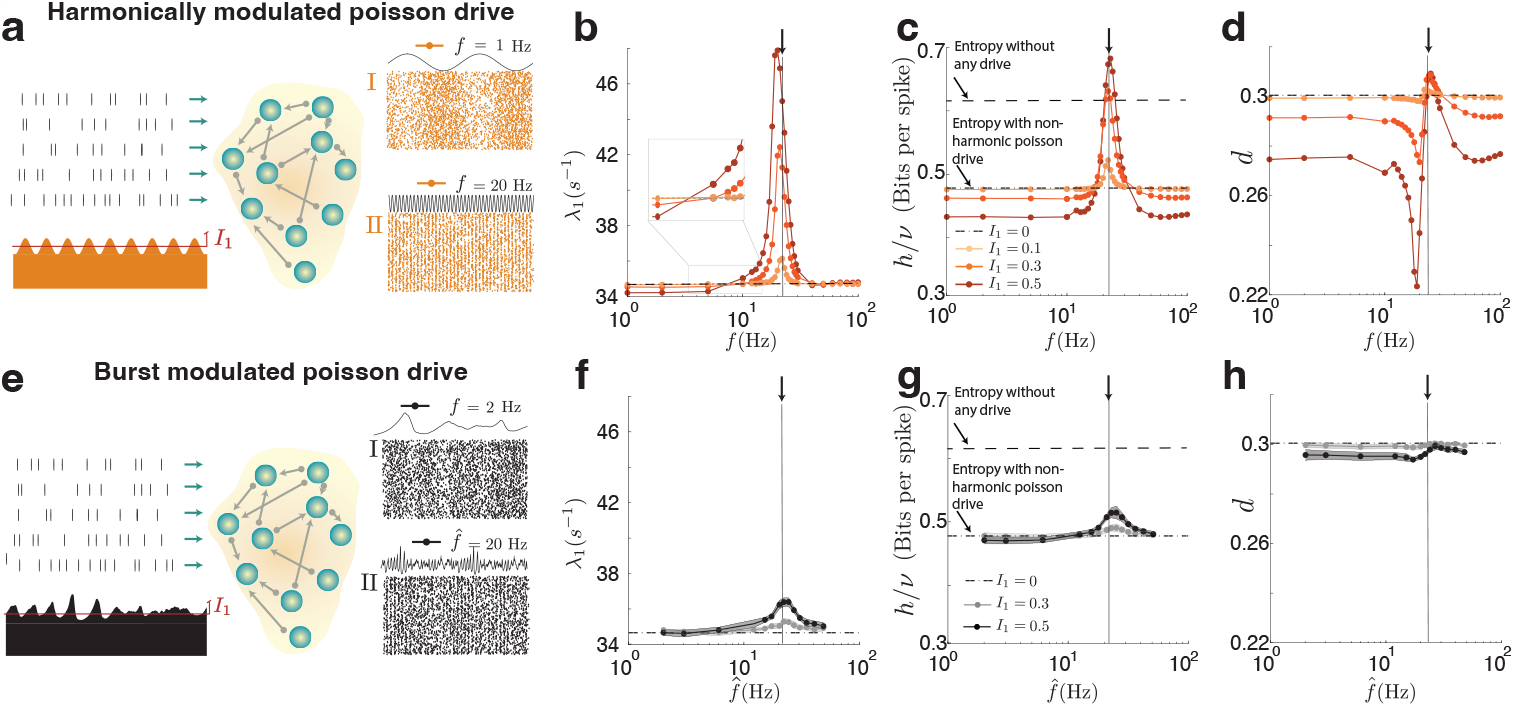
Resonant boosting of chaos is quenched by transient synchrony. **(a)** (left) Diagram of a non-delayed inhibitory network driven by harmonic external input. Each neuron receives, besides its recurrent input, K external spike trains sampled from an inhomogeneous Poisson process with a harmonically modulated rate with baseline *I*_0_, amplitude *I*_1_, and frequency *f*, designed to mimic the oscillatory input after the transition to oscillations in Figure 3 **(a)** (right) Raster plots of a network activity driven by external inputs of frequencies *f* = {1, 20} Hz. **(b)** maximum Lyapunov exponent, **(c)** metric entropy and **(d)** attractor dimension, as a function of the frequency of the cosine-modulated Poisson input *f*, for increasing values of the input drive’s oscillatory amplitude *I*_1_. The top dashed line in panel (c) is the value the entropy with a constant input (not Poisson, i.e. that of Fig. 3). The bottom dashed line is that of a non-harmonically modulated Poisson drive. Notice how the external input drive reduces chaos proportionally to the intensity of the drive for smaller frequencies and how this behavior is reversed in the neighborhood of the resonant frequency of the network.**(e)** (left) Diagram of a non-delayed inhibitory network driven by an external input with transient oscillations. Each neuron receives K external spike trains sampled from an inhomogeneous Poisson process with a rate given by transient oscillations of a different spiking network model (see (10) and *Methods*) with baseline *I*_0_, amplitude *I*_1_, and peak frequency 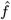. **(e)** (right) Raster plots of a network activity driven by external inputs of with peak frequencies of 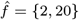 Hz. **(f)** Maximum Lyapunov exponent, **(g)** metric entropy and **(h)** Attractor dimension, as a function of the frequency of the transient-oscillation-modulated Poisson input 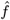, for increasing values of the input drive’s oscillatory amplitude *I*_1_. Notice how, compared to the harmonic case, a drive with transient synchrony and drifting frequencies, recruits responses of the network that would either reduce or intensify chaos.

The network has a non-monotonic response to oscillatory inputs of increasing frequency, as indicated by the pronounced peak of synchronization index in Fig. **??b**. If we were to increase the coupling strength of the network to a critical value of K, this peak would develop into a full resonance (38). The generation of self-sustained oscillations at the population level shown in Fig. 3 is reached when the drive that generates a resonance in the network is internally generated. Here we show that for values of weaker coupling than the critical ones, driving the network with an oscillatory drive already shows signatures of this developing resonance, and that being in the neighborhood of that resonance is a sufficient condition for chaos to be intensified.

The network response to the weakest amplitude of the modulation (peach line, *I*_1_ = 0.1), is the scenario that better captures the activity of the network at the onset of the oscillatory instability in the non-driven network. In that case, the maximum Lyapunov exponent (Fig. 6**b**), the metric entropy (Fig. 6**c**) and the attractor dimension (Fig. 6**d**) all peak at a drive frequency of 21 Hz (see black guiding line).

The values of the frequency of the external input at which the maximum Lyapunov exponent, the metric entropy and the attractor dimension change from being suppressed by the external drive to being boosted by the drive are different for each measure. This can be better understood by looking at how the entire Lyapunov spectrum depends on the driver frequency (Supplementary Fig. S12**c**). With increasing frequency of the external drive, the positive Lyapunov exponents become more positive, but the negative ones become more negative at earlier frequencies and grow more strongly than the positive ones, resulting in a sharp decrease of the attractor dimension (fewer negative exponents are needed to cancel the sum that defines the dimension, see Fig.2**f**).

Oscillatory activity in the cortex is not perfectly periodic, it is usually confined to short episodes with frequencies that fluctuate over time (53). This type of transient oscillatory activity can be leveraged to flexibly gate information between areas (10), but whether a network driven by such transient oscillatory activity can enhance or reduce network chaos remains unknown. Here we asked whether driving a network with transient oscillatory bursts would lead to an increase in noise entropy, or if the broad spectrum of these signals would allow recruitment of suppressive responses, resulting either in weak modulation or overall suppression. We generated an inhomogeneous Poisson process with a rate modulated by transient oscillatory bursts, which were simulated as described in (10), and drove a non-delayed network with this drive (Fig. 6**d**, left). We changed the peak frequency of the signals 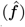 by re-scaling the temporal axis (see *Methods*). We found that this type of drive largely cancels the intensification of chaos at the resonant frequency. The dependence of the maximum Lyapunov exponent on the peak frequency of the oscillatory burst of the input drive 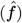 is shown in Figure 6**fii**. The maximum Lyapunov exponent is only significantly modulated in the neighborhood of the resonant frequency, having only a 5% increase rather than the almost 40% increase in the harmonic drive case. The metric entropy is also only weakly modulated by this type of drive. Because the bursts are transient and the frequency of the burst is not fixed, this input drive recruits suppressive responses from low frequencies, leading to an overall weak modulation of the metric entropy (Fig. 6**g**) and a reduction of the attractor dimension (Fig. 6**h**).

## Discussion

### Delayed interactions in dynamical systems

We developed a framework that allowed us to compute, for the first time, the entire spectrum of Lyapunov exponents in an effectively delayed high-dimensional system and therefore exactly characterize the dynamics on the attractor. We note that delayed systems can have an infinite-dimensional phase-space exploration, and therefore a *discrete* spectrum of Lyapunov exponents is not guaranteed to exist. We mapped the pulse-coupled spiking network with delayed interactions to an equivalent system with fixed and finite degrees of freedom. We found that the Lyapunov spectrum is invariant to network size, giving rise to extensive chaos, with a finite dimension of the attractor for finite network size (Fig. 2), as also found in other systems with infinite-dimensional phase space (54). We showed that, prior to the oscillatory instability, the attractor dimension increases linearly with the interaction delay, while the top 5% of the LEs (including the maximum LE) decrease in magnitude, while the number of positive LEs increases. This tension between a reduction in magnitude and an increase in the number of LEs leads to the remarkable fact that the metric entropy of the network is independent of the delay up to the oscillatory instability (Fig. 3). These same exact dependencies were found in classic studies of delayed dynamical systems, for either low-dimensional maps or scalar differential equations (28, 29, 35, 36). Remarkably, we find that these dependencies are also present in weakly delayed rate networks (Figs. S12-S13), where the transition to oscillations is accompanied by a complete collapse of chaos. These findings suggest that, in the absence of a mean-field bifurcation, the reshaping of the attractor dynamics with increasing delay is of a universal nature.

### Chaos in spiking networks

Balanced networks, as originally described for binary neurons, are chaotic with an infinitely large maximum Lyapunov exponent (39). Although the statistics of the balanced-state networks appear to be largely independent of the single neuron model (3, 39, 55), its dynamical stability is not common or intrinsic to the balanced state. Random networks of pulse-coupled, inhibitory leaky integrateand-fire (LIF) neurons have stable dynamics (55, 56): After a transient of irregular activity whose duration scales exponentially with network size, the dynamics settle in a periodic orbit (56, 57). However, finite size perturbations reveal a phase space of simultaneously diverging and contracting trajectories (58, 59). Trajectories initially separated on average by less than a critical value, converge to each other exponentially (local stability), while those separated further diverge exponentially. The inclusion of longer time scales, via synaptic dynamics, can lead to a loss of dynamical stability, resulting in extensive chaotic dynamics after a critical value of the synaptic time constant (56, 60, 61).

Incorporating a dynamical mechanism for the generation of the action potential, as studied here for the quadratic integrate-and-fire neuron, leads to extensive chaos to sufficiently large external input (3, 6) even in purely inhibitory pulse-coupled networks. Here, we showed that the inclusion of delays does not affect the extensive nature of chaos, as including delays does not lead to the development of deterministic chaos in LIF neurons (56). We have not observed any transition to dynamical stability (i.e. to a negative maximum lyapunov exponent) that coexists with spiking variability, and chaos only collapses when the dynamics converge to clock-like synchrony. Future work will need to determine whether oscillatory activity can actively modulate the number of neurons contributing to the most stable directions at any given point in time, given that those neurons can respond reliably to the same presentation of the stimulus despite the chaotic dynamics (5, 6)

### Collective oscillations and chaos

Our results demonstrate that the emergence of sparse, collective oscillations induces a complete dynamical reconfiguration in models of the cortical circuitry. We find that increasing the recurrent delayed inhibition, either by increasing the synaptic delay *δ* (Figs. 3-5), or the number of presynaptic inhibitory inputs (Supplementary Fig. S6), leads to a transition from an asynchronous irregular state characteristic of balanced-state networks, to a state with irregular activity at the single cell level but with collective rhythms at the populations level. This transition occurs via a Hopf bifurcation, analogous to that described previously in models lacking a dynamical mechanism for the generation of the action potential (15, 16, 31). Recent theoretical work has described the transition to collective oscillations of the QIF with increasing in-degree (62, 63), and work has also described the mean field dynamics of similar networks to us, reporting rate chaos (chaos in the mean field equations) with increasing input current to the cells (64). This is very different from the chaos reported here, in which the population rates are not chaotic. Previous work has found a coexistence of chaotic dynamics with sparse oscillations in networks of LIF neurons with slow synapses (4), but how oscillatory activity impacts the chaotic dynamics of the network remained elusive.

We used a powerful approach (38) to compare the equivalent system studied here and the true, infinite-dimensional delayed system and found that at the onset of the theoretically predicted value of the synaptic delay for the onset of collective oscillations (Fig. 4) chaos is greatly intensified, by simultaneously increasing the maximum Lyapunov exponent, the dimension of the chaotic attractor and the metric entropy. This finding is insensitive to the presence of single-cell (Figs. 5, S8, S9) and cell-type (5, S11) heterogeneity, and holds true in the entire region of the parameter space analyzed (S10). We found that this reconfiguration is only possible in models that explicitly incorporate a dynamical mechanism for the generation of the action potential and is not present in rate models (Figs. S12 and S13), in which we have not found chaotic, asynchronous activity to coexist with collective oscillations (but see (65) for oscillatory rate networks in the presence of structured connectivity).

To investigate the mechanisms underlying such intensification of chaos, we studied a non-delayed network driven by a harmonically-modulated Poisson process. We found that chaos is reduced for low frequencies of the drive, analogously to what is found in noise-driven rate networks (66). Nevertheless, for frequencies in the neighborhood of the resonant frequency, the chaotic measures are intensified. This boosting of chaos with an external asynchronous oscillatory drive is to our knowledge one of a kind and has strong resemblance to what is found in classic maps driven by a harmonic input. This finding is in stark contrast to what is found in rate network models (14, 50), for which a sufficiently strong harmonic drive is able to completely abolish chaos at all frequencies of the harmonic drive.

### Transient oscillations lead to moderate boost of network chaos

Oscillatory activity in cortex is far from perfectly periodic, cortical rhythms are in fact characterized by transient oscillatory bursts of drifting frequency (53, 67) that occur in a seemingly stochastic fashion. Theoretical work has shown how to flexibly route information, leveraging those noise-induced transients of rhythmic activity (10). Our results here show that when input spike trains of the network are modulated by a transient oscillatory amplitude, the response of the network to such input, recruits suppressive responses from lower frequencies, leading to a very modest change in the Lyapunov exponent and in the metric entropy, and a mild decrease in the attractor dimension. These effects are in strong contrast with the periodic drive case studied, in which the maximum Lyapunov exponent and the metric entropy undergo a 40 % increase compared to the non-driven case. Future work will need to elucidate whether input patterns generated by a chaotic network with transient oscillatory burst would lead to chaos suppression as seen here with Poisson spiking patterns, or would lead to an increase of chaos in the target network.

We emphasize that generally, in dissipative systems (including our networks and possibly in contrast with Hamiltonian models (68–70)) a positive entropy rate is possible despite the existence of an invariant measure (71) (i.e. despite the fact that probability density over the attractor is not changing over time), because the measure is singular and therefore is a “bottomless source of entropy” (72, 73).

## Materials and Methods

### Network Model

We considered a network of N quadratic integrate-and-fire (QIF) neurons

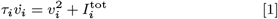

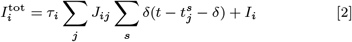

where *τ*_*i*_ is the membrane time and *v*_*i*_ is the voltage of the neuron *i*. The term 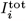 describes the synaptic current that a neuron *i* receives at time *t* given that its pre-synaptic neighbors *j* emitted spikes in the times 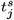 after a delay *δ*. The connectivity *J*_*ij*_ has a binomial distribution with probability *K/N* where 1 ≪ *K* ≪ *N*. When a connection exists, its strength takes a constant value depending on the type of the pre and post cell: 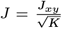, for *x, y* = *E, I*. The second term is an constant external input 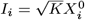, *X*^0^ = *E*_0_, *I*_0_ for excitatory and inhibitory neurons respectively (see Section A in *Supplementary Information*)

#### A. Phase reduction for integrate-and-fire models

In networks of pulse-coupled units receiving constant external inputs, and whose voltage dynamics have a defined threshold *x*_*t*_ and a reset values *x*_*r*_, the evolution between network spikes *t*_*s*_ is defined by a propagator function or map (25, 37, 74–76)

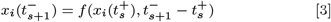

The function *f* evolves the state of the neuron *i* after the last spike in the network, 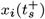, to the state just before the next spike at *t*_*s*+1_. If the neuron *i* is not postsynaptic to the neuron *j*^*∗*^ that produces an spike at *t*_*s*+1_, the state variable will be left unaffected, and then 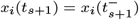. On the contrary, if the neuron *i*^*∗*^ is postsynaptic to *j*^*∗*^, then its state variable has to be further updated:

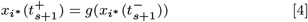

Eqs 3 and 4, form the basis of the iterative evolution of the network from spike to spike. The Eqs for the QIF neuron used in this paper were derived in Ref. (3), a detailed derivation can also be found in Section B of the *Supplementary Information*)

#### B. Delayer single-compartment-axon

Synaptic and axonic delays consistent with the framework above specified were incorporated into the network by means of the introduction of *delayer* variables, chosen to be QIF neurons in the excitable (non-periodically spiking) regime.

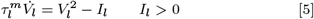

The solution of Eq. 5 for all three cases (i) 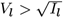, (ii)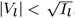, (iii) 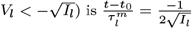 ln 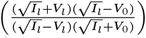. The transition between the regimes (i),(ii),(iii) is dynamically forbidden.

We can define a change of variables

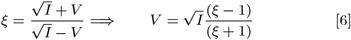

that simplifies the solution of Eq. 5 reads:

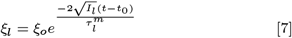

In this representation a propagator function *η* and an update function *γ*, analogous to *f* and *g* of Eqs. (3,4) can as well be defined:

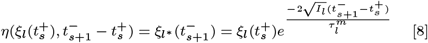

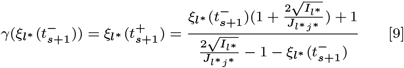

The *ξ* variable, when initially below the threshold value *ξ*_*t*_ = −1 *ξ*_0_ =*<* −1 takes a finite time to spike given by 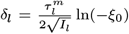 (see Figure S1). If the reset value is set to *ξ*_*r*_ = 0, there will be no dynamics after the spiking event. If while in its resting state, an incoming spike from neuron *j*^*∗*^ is received, then its state will be modified according to *γ*(0) and will depend only on the parameters of the system 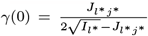. If the incoming spike has a sufficiently large weight then the *ξ* variable will be pushed beyond its unstable fixed point and emit a spike after a fraction of time given by:

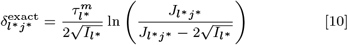

before relaying it to the next neuron. More concretely, the value of *J*_*l∗j∗*_ has to be bigger than 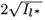, the distance between the fixed points in equation 5. For the case in which *J*_*l∗j∗*_ is independent of the neuronal pair, the delay introduced by the SCA will only depend on its own parameters, and then *δ*_*l∗j∗*_ = *δ*_*l∗*_. The singlecompartment-axon (SCA), is then defined by Eqs. 9, with a reset value *ξ*_*r*_ = 0 for the “exact delay” framework. If the SCA has a reset value that is equal to any value between the threshold *ξ*_*T*_ = −1 and zero, the delay will depend on the dynamics of the network and change from spike to spike. The delay, in that case, will be :

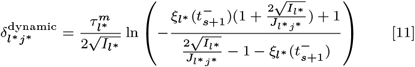

For *J >* 0, this equation has solutions for 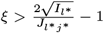. The smaller the value of 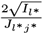, the larger the range of *ξ* to which it can be reset, and the faster the function in the logarithm reaches a value that is only weakly dependent on *ξ*. This configuration is what we will can call “dynamic delay” framework.

In both cases, when the SCA is arranged post-synaptically to each neuron in a network, it will delay the transmission of the spike by an amount given by Eq. 10 or 11, while preserving the desirable features of iterable maps.

#### C. Equivalent delayed network

A diagram of the network architecture of a delayed network with balanced state properties is shown in Fig. 2**a**. A spiking neuron *j*^*∗*^ (in orange, left), described by its phase variable *ϕ*_*j*_ receives independent and large inputs proportional to the square root of the average amount of connections per neuron *K*. Its spike is instantaneously transmitted to the SCA, of variable *ξ*_*l*_ in green. The parameters of the SCA define the time it will take for the spike to be transmitted, from the SCA to the postsynaptic neurons *i*, whose phase variable is *ϕ*_*i*_.

The connectivity matrix, 𝕁 has then the form in Eq. 12. For ease of notation, indexes *i* and *j* (from 1 to N) will be reserved for phase neurons and *l* and *m* (from N+1 to M = 2N) for the SCA. The connectivity matrix is a block matrix describing an ErdősRényi random graph on the upper right block, with weights proportional to 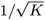 and a diagonal matrix in the bottom left:

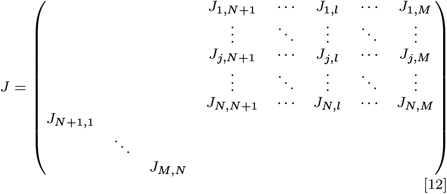

The event-based simulation of the system defined in Eqs (1, 2) is done by, after each spike, evolving the network dynamics by propagating both the somas by Eqs. S10 and the SCA 9 up to the next spike time. If the next spike time is from a soma, then the postsynaptic SCAs have to be updated according to Eq. 9, and if it was from an SCA, postsynaptic somas should be updated as in Eq. S10.

#### D. Characterization of network activity

The firing rates of each neuron *ν*_*i*_ were calculated by summing all the spikes and dividing by the time spanned between the first and the last spike. The mean population rate is then *ν*. The coefficient of variation is the square root of the squared inter-spike-interval times the squared rate, i.e. 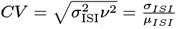. The level of synchrony of the network was assessed by the synchronization index 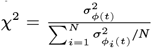 Where, 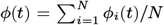 is the mean activity in the phase representation. The variables *σ*_*ϕ*(*t*)_ and *σ*_*ϕ* (*t*)_ are the standard deviation of the mean phase over time or, respectively, of the phase trace *V*_*i*_(*t*) of each individual neuron *i*. The *χ* coefficient is bounded to the unit interval 0 *< χ <* 1, with vanishing values indicating asynchronous dynamics. The amplitude *A* (Hz) and the frequency *f* (Hz) of the population activity were calculated by making a binning histogram of a 1 ms bin of the spikes of the entire network. This quantity normalized by the bin size and the number of neurons defines a multi-unit like signal (MUA). The mean peak high defines the amplitude *A* (Hz). The frequency of the MUA was obtained as the inverse of the first autocorrelation peak.

#### E. Simulation specifics and parameters

The simulations took default parameters given by table 1. Deviations from these default parameters are indicated below. The simulations are event-based using the iterative map described in the previous sections and following Ref.(3). In short, one needs to calculate the next spike time in the network, coming either from a soma or from a SCA. For that, it is sufficient to find the soma with the largest phase and the SCA with value closer to 1. For each simulation, random topologies (ErdsRényi) for the neuronal block (Eq. 12), random initial conditions, and an initial guess on the input currents to the neurons are generated. By means of root finding algorithms (Regula Falsi and Ridders’ method) (77), a guess on the input current *I* needed to have a target firing rate was made from Eq. S3. After a simulation lasting S_R_ spikes per neuron, the rate was calculated and a new guess on the currents is made. This procedure is then iterated until the target mean firing rate is found with 1% precision. The multiplying factor that measures the distance to the balance condition is defined by *Is* = *I/I*_0_. Once the appropriate value of the current *I* is found, the network can be warmed up to disregard transients, which can be large in delayed systems by a time equivalent to S_W_ spikes per neuron. Finally, a random orthonormal matrix Q is chosen and the QR algorithm described in the theory section below was left to warm up for a time duration equivalent to S_WONS_ spikes. This guarantees some degree of alignment of the first Lyapunov vector to the first vector of the orthonormal system. The simulation runs for a time equivalent to S_C_ spikes per neuron.

**Table 1.**
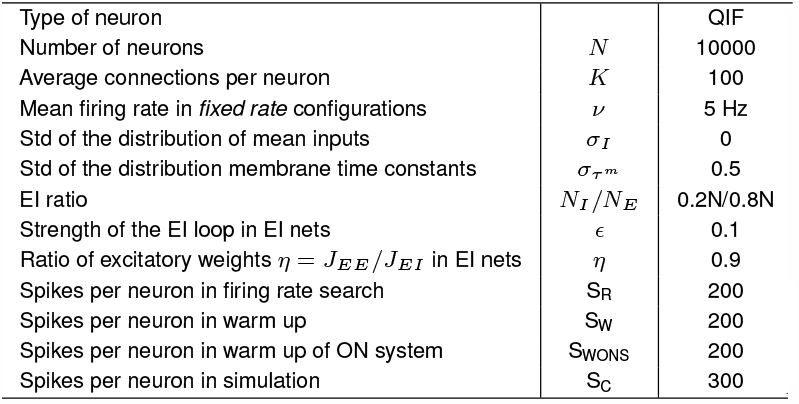
List of default network parameters.

|  |  |  |
| --- | --- | --- |
| Type of neuron |  | QIF |
| Number of neurons | $N$ | 10000 |
| Average connections per neuron | $K$ | 100 |
| Mean firing rate in *fixed rate* configurations | $\nu$ | 5 Hz |
| Std of the distribution of mean inputs | $\sigma_I$ | 0 |
| Std of the distribution membrane time constants | $\sigma_{\tau^m}$ | 0.5 |
| EI ratio | $N_I/N_E$ | 0.2N/0.8N |
| Strength of the EI loop in EI nets | $\epsilon$ | 0.1 |
| Ratio of excitatory weights $\eta = J_{EE}/J_{EI}$ in EI nets | $\eta$ | 0.9 |
| Spikes per neuron in firing rate search | $S_R$ | 200 |
| Spikes per neuron in warm up | $S_W$ | 200 |
| Spikes per neuron in warm up of ON system | $S_{WONS}$ | 200 |
| Spikes per neuron in simulation | $S_C$ | 300 |

Deviations from the default which are not indicated explicitly are as follows: In Figure 5 N=8000 for panels **f-j**. Figure S2 had N=400, while in S10(g-h),(p-q), (y-z) N was 2000, and in those cases S_R_=S_Q_=S_WONS_=600. In Figure S10 N=5000.

In Figures 3, 4, S3, S4, S6, S7 no heterogeneity was present (i.e.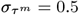).

The driven network simulations of Fig. 6 were performed as follows: Each neuron in the recurrent network receives an independent stream of inhibitory external input spikes of strength 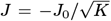 that are generated from an inhomogeneous Poisson process with a rate that is modulated either I) sinusoidally with frequency *f* and amplitude *I*_1_ around a constant rate *ν*_in_ = *Kν* or II) by the outcome of the LFP obtained by simulations of networks with transient synchrony (identical to that in Fig 1 of (10)). We implemented the sinusoidally modulated input drive for each neuron by drawing the inter-spike intervals of the inhomogeneous Poisson input from the distribution *p*(Δ*t*_ISI_) = − log(*x*)*/*(*ν*_in_*τ* ^m^(1 + *I*_1_ sin(2*πf t*)), where *x* is uniformly distributed between 0 and 1, *I*_1_ is the relative amplitude of the input modulation, *f* is the frequency of the input modulation and *ν*_in_ is the external input rate that we chose to be 5 times the mean firing rate *ν*. We set *J*_0_ = 1. This implementation results precisely in the desired sinusoidally modulated Poisson rate for a relative input modulation strength *I*_1_ ≪ 1 and *f* ≪ *ν*_in_. By adapting the constant external input current iteratively using the bisection scheme described above, we obtained a target firing rate of 5 Hz. We chose *ν*_in_ = *Kν* = 500Hz, thus that on average neurons in the recurrent network receive the same number of recurrent spikes as external input spikes. After a simulation lasting S_R_ = 10^3^ spikes per recurrent neuron, the rate was calculated and a new guess of the currents was made. This procedure is iterated until the target mean firing rate was found with 1% precision. Once the appropriate value of the current *I* is found, the orthonormalization procedure described below was done iteratively where the interval size was adapted to achieve a condition number of the *Q* matrix between 1e2 and 1e6 with an initial guess of 5000 spikes. We performed a warmup of both the recurrent network state and the state of the orthonormal system. The simulation ran for a 60s. Results in Fig. 6 show averages over 10 ErdsRényi network realizations.

#### F. Calculation of the critical delay

In the regime in which each neuron receives a large number of inputs per integration time, with each input making only a small contribution to the net input, and under the assumption that all neurons are the same, the dynamics of the network can be analyzed under the so-called *diffusion approximation* (15, 31, 38, 78). We used the method described in (38) to do this calculation, and is detailed in the Section C in the *Supplementary Information*

#### G. Ergodic theory measures characterizing the attractor

The dimensionality *D* of the attractor of the delayed circuit can be then computed by means of a classic conjecture by Kaplan and York (79). The conjecture states that the attractor dimension is given by the linear interpolation of the number of Lyapunov exponents that are needed for its cumulative sum to vanish (see Fig. 2**f**, top, see also Sect. A in *Supplementary Information*):

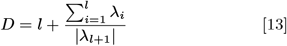

Where *l* is such that 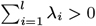 and 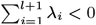.

The metric entropy *H*, is an upper bound on the rate at which the chaotic dynamics contributes to the Kolmogorov-Sinai entropy, in other words, an upper bound on the average information gained by making a new measurement in a sequence of measurements of the state of the dynamical system (32) (see also Sect. A of *Supplementary Information*). We compute this metric entropy via the Pesin identity (80), which equates the metric entropy with the sum of the positive Lyapunov exponents (see also Fig. 2**f**):

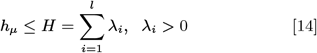

In Figs. 3,5 and 6, we show the intrinsic metric entropy in bits per second, normalized by the mean firing rate of the network. In other words

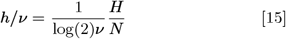

Keeping in mind that the measures derived from the Lyapunov spectrum are upper bounds and that no rigorous statement can be made in favor of the equalities, we nevertheless refer to them as the metric entropy and the attractor dimension.

#### H. Lyapunov spectrum of delayed spiking neuronal networks

Given a neuronal model *F* (*V*_*i*_) and a model for the SCA that are exactly solvable, the network can be propagated between spikes and updated by Eqs. S10 and S7. From these equations, a Jacobian 𝕃 (***x***(*n*)) at each spike time can be obtained, from which the estimation of the Lyapunov dimension *D* (Eq.13) and the metric entropy *H* (Eq.15) are possible via QR decomposition. The terms of the Jacobian and an explanation of the terms can be found in Section B of the *Supplementary Information*.

## ACKNOWLEDGMENTS

We thank M. Monteforte and M. Puelma Touzel for discussions, and L. Abbott and T. Nguyen for feedback on this manuscript. This work was supported by the German Research Foundation (Deutsche Forschungsgemeinschaft, DFG) through SFB 1528, SFB 889, SFB 1286, SPP 2205, DFG 436260547 in relation to NeuroNex (National Science Foundation 2015276) & under Germanys Excellence Strategy - EXC 2067/1-390729940; by the Leibniz Association (project K265/2019); and the Niedersächsisches Vorab of the VolkswagenStiftung through the Göttingen Campus Institute for Dynamics of Biological Networks.This work was partially supported by the Federal Ministry for Education and Research (BMBF) under grant no. 01GQ1005B (to A.P., R.E. F.W.), by a GGNB Excellence Stipend of the University of Göttingen (to A.P.), by a Swartz Fellowship for Theory in Neuroscience (A.P.) through CRC 889 by the Deutsche Forschungsgemeinschaft and by the VolkswagenStiftung under grant no. ZN2632 (to F.W.)

## Supporting Information Text

### 1. Delayed spiking networks

Here we describe the methods for Figs. 1-7. All spiking network simulations were simulated in an event based manner as described in the sections below.

#### A. Network Model

We considered a network of N quadratic integrate and fire (QIF) neurons

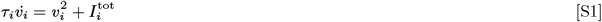

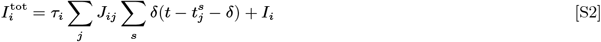

where *τ*_*i*_ is the membrane time and *v*_*i*_ is the voltage of the neuron *i*. The term 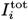 is the total input current, consisting on two terms. The first one describes the synaptic current that a neuron *i* receives at time *t* given that its pre-synaptic neighbors *j* emitted spikes in the times 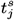. Every time a neuron spikes, takes a time *δ* to arrive to its post-synaptic target. The connectivity *J*_*ij*_ has a binomial distribution with probability *K/N* where 1 ≪ *K* ≪ *N*. When a connection exists, its strength takes a constant value that only depends on whether the presynaptic and postsynaptic cells are excitatory or inhibitory.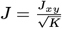, for *x, y* = *E, I*. The second term is an constant external input 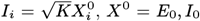, *I*_0_ for excitatory and inhibitory neurons respectively.

The mean firing rate of the E and I populations is given by (1, 2) 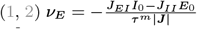 and 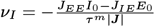. The mean input current is given by 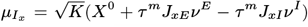; where *X*_0_, is a positive (excitatory) input current to the population *x*: *E*_0_ for the excitatory population and *I*_0_ for the inhibitory one.The variance of the input currents are given by 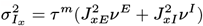

For the particular case of networks of inhibitory neurons, the condition of a finite mean input current requires that 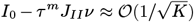. In the limit of large K the firing rate must then satisfy 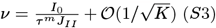. For networks of excitatory and inhibitory neurons, we request that the first moment of the firing rates distributions are the same (*ν*_*E*_ = *ν*_*I*_), and that the variances of the input currents are equal 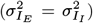. Together, these conditions impose further constraints on the synaptic weights: 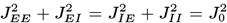. The weight matrix is written as a function of three parameters: *J*_0_ *η* = *J /J <* 1 and *ϵ* = *J /J* (5). The synaptic weights then have the form 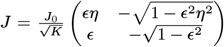. The stable solutions with positive rates are found for *η* and *ϵ* such that 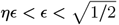 (5).

#### B. Phase reduction for integrate and fire models

In networks of pulse coupled units receiving constant external inputs, and whose voltage dynamics have a defined threshold *x*_*t*_ and a reset values *x*_*r*_, the evolution between network spikes *t*_*s*_ is defined by a propagator function or map (6–10)

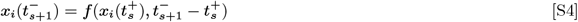

The function *f* evolves the state of the neuron *i* after the last spike in the network, 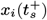, to the state just before the next spike at *t*_*s*+1_. If the neuron *i* is not postsynaptic to the neuron *j*^*∗*^ that produces an spike at *t*_*s*+1_, the state variable will be left unaffected, and then 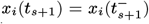. On the contrary, if the neuron *i*^*∗*^ is postsynaptic to *j*^*∗*^, then its state variable has to be further updated:

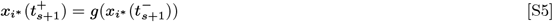

Eqs, S4 and S5, form the basis of the iterative evolution of the network from spike to spike. Its precise form will depend on the neuron model considered, as will the expression for the time to the next spike.

Analytically solvable neuron models which in absence of any recurrent input have periodic spiking activity can be mapped to a phase variable that linearly evolves in time. This variable, *ϕ*, has a propagator function *f* (*ϕ*) that is a linear function of the inter-spike times. All the information about the particular neuron model can be condensed in the form of the update function *g*.

The QIF neuron defined in Eq. S1, is analytically solvable in either its supra-threshold or its excitable regime; different solutions of Eq. S1 can be obtained depending on the sign of *I*_*i*_ (in the absence of recurrent connections). In the supra-threshold regime (*I >* 0) the solution to Eq. S1 with initial conditions *V* (*t*_0_) = *V*_0_ is

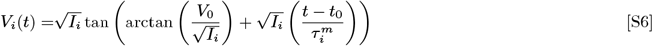

The propagator function *f* and the update function *g* in the voltage representation for the supra-threshold QIF are then:

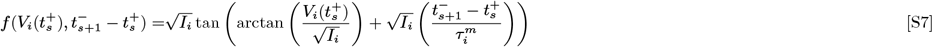

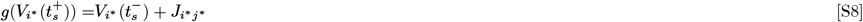

We see that from Eq. S7, that the variable change *ϕ* = 2 arctan 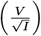 reduces the dynamics to a linearly evolving phase, *ϕ* ∈ (−*π, π*). The free period can be obtained requesting that *V*_*t*_ = +∞ and *V*_*r*_ = −∞ : 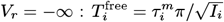. This yields a value for the frequency 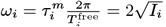. The propagator and update functions for *ϕ* are

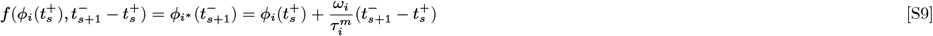

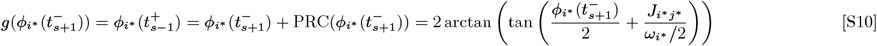

Where PRC is the phase response curve.

#### C. Calculation of the critical delay

Here we described the method to describe the transition to population oscillations and how to obtain the critical delay following the approach by Richardson (11, 12). The single neuron dynamics are then given by 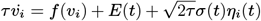. Here, *E*(*t*) = *τ KJν*(*t − δ*) and 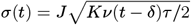 calculated in standard manner under the assumption of uncorrelated activity in the asynchronous irregular state (2–4), *f* (*v*) = *v*^2^ for the case of the QIF neuron here considered and *η*_*i*_(*t*) is white gaussian noise. If the number of inputs per neuron *K* is fixed, then the dynamics of the distribution of voltages *P* (*v, t*) are described by a Fokker-Plank equation.

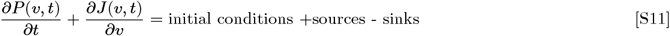

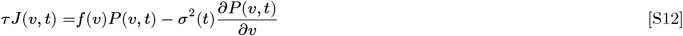

where *J*(*v, t*) is the probability current and the voltage-independent diffusion coefficient is given by 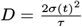. Richardson (11, 12) developed a numerical method to i) evaluate numerically the steady state solution *p*_0_(*v*) and find self consistently the steady state firing rate 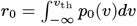 and ii) the linear response to parameter modulations, which allows to self-consistently calculate the critical parameters where the asynchronous irregular state loses stability to a synchronous irregular one. Here we give a brief summary of how to compute, for an otherwise fixed set of parameters, the critical delay *δ*_*c*_. If *E* and *σ* are constant, a steady state will be reached in which the probability distribution is independent of time. In this case, the stationary firing rate can be self consistently found by the normalization condition of the probability distribution. This was done analytically in (3, 4) for the leaky integrate and fire (LIF) neuron and in (13) for the quadratic integrate and fire (QIF). Although the general-time dependent solution for nonlinear equations is inaccessible, significant progress can be made by studying the linear response to a harmonically modulated parameter. *α* = *α*_0_ + *α*_1_ exp(*iωt*).

We first solve the harmonic modulation of the input current *E*(*t*) for a network of uncoupled neurons, following Ref (11) and then ask for self-consistency. To do this, we decompose the probability current 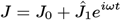, the density 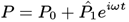, the input current *E* = *E*_0_ + *E*_1_*e*^*iωt*^ and the firing rate 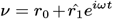. Inserting those in Eq. S12, and separating contributions to the modulation coming from *E*_1_ and from 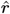 (see sections 3.3 and 4.1 in Ref (11) for details) we obtain two equations. One, defines the *E*_0_ that satisfies the steady state with the target firing rate *r*_0_. The other is the equation that relates 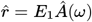, where *Â*(*ω*) is a function of the frequency *ω*, obtained using theRef (11) method.

### Calculation of the critical delay

To compute the critical delay that destabilizes the asynchronous irregular state, we need to require that the input modulation *E*_1_, which had so far been considered external to the network, arises from network activity. This means that *E*(*t*) = *τ KJν*(*t* − *δ*) ≈ *E*_0_*r*_0_ + *E*_1_*e*^*iωt*^, 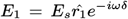 where *E*_*s*_ = *τ KJ <* 0 (for inhibitory networks). The self-consistent equation (analogous to Eq. (52) in Ref (11), reduced to the delta coupled network case) is 1 = − |*E*_*s*_| *e*^−*iωδ*^*Â*(*iω*). Writing the (known function) *Â* = |*A*(*ω*)| *e*^*iϕ*(*ω*)^, separating in real and imaginary part we have effectively two equations with two unknowns:

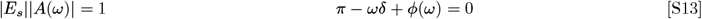

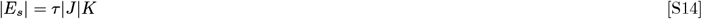

From which we can obtain the value of the frequency *ω* and the critical delay *δ* that destabilizes the asynchronous irregular state.

### 2. Ergodic theory of effectively delayed spiking neuronal networks

#### A. Background and definitions

The simplest measure of the dimensionality of a chaotic attractor is called Kolmogorov capacity or box-counting dimension (14). It is a metric-based measure in the sense that it does not take into account the density of orbits on the attractor. Given a parallelepiped of side *ϵ*, the number *N* (*ϵ*) that is needed to cover the set of points conforming the attractor *A* is expected to satisfy *N* (*ϵ*) ≈ *ϵ*^−*dc*^. The capacity of the set can then be defined as 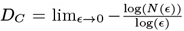. If instead of considering the number of hypercubes of side *ϵ* to cover the attractor, we consider the number of hypercubes *N* (*ϵ, ϑ*) needed to cover a fraction *ϑ* of the attractor, then the *ϑ* capacity can be defined as 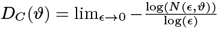. This measure, which for *ϑ* = 1 is directly the capacity and otherwise will be called *D*_*µ*_, satisfies *D*_*µ*_ ≤ *D*_*C*_. Finally, a point-wise dimension *D*_*p*_(*x*) can be as well defined by estimating the exponent with which the total probability within a ball of radius *ϵ* decreases as the radius vanishes (14) 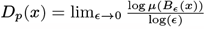.

For a definition of an attractor dimension that is related to the measure of the attractor, it is necessary to define a partition ℬ= {*B*_*i*_} that covers the phase space. At each measurement, the trajectory of the system can be found in one *B*_*i*_ such that for sufficiently long times, a frequency of occurrence *P*_*i*_ can be assigned to each *B*_*i*_. When the size of the elements of the partition tends to zero, *P* defines a probability density such that its sum over the attractor is equivalent to the measure on the attractor *µ*(*A*). Then, the frequency of occurrence can be written as 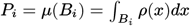.

When observing the state of the dynamical system at a given point if time, the information that is gained about the state depends on the resolution of the instrument, the *diameter* of the partition used. This diameter is basically given by the largest size of the elements *B*_*i*_ of the chosen partition. If the partition has diameter *ϵ*, then the information gained by making a measurement is given by *I*(ℬ (*ϵ*)) = − ∑_*i*=1_ *µ*(*B*_*i*_(*ϵ*)) log (*µ*(*B*_*i*_(*ϵ*))). If now, from all the possible partitions to be chosen, we choose that one that minimizes the expression, then the information dimension *D*_*I*_ is defined as 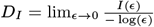.

Kaplan (15), defined a measure of an attractor, the Lyapunov dimension, as a function of the Lyapunov exponents:

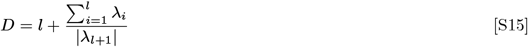

Where *l* is such that 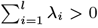 and 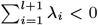. Kaplan (15) conjectured initially that the Lyapunov dimension *D* was generally equal to the fractal (box counting) dimension, or a lower bound to it (14). This form of the Kaplan-York conjecture can be found in the literature (15, 16) and in text books (17, 18). In further work by both Kaplan and York (19) and York and colleagues (14), this conjecture was updated to a form which includes information of the density of orbits over the attractor. They specifically conjecture that *D* = *D*_*C*_ (*ϑ*). as in Ref. (14) further conjectured that this equality and the original Kaplan York conjecture all hold. Rigorous results show that *D*_*P*_ = *D*_*I*_ = lim_*ϑ*→1_ *D*_*C*_(*ϑ*) ≤ *D* for invertible smooth maps (as reviewed in (14, 20)). The conjecture has been shown to hold whenever a Sinai-Ruelle-Bowen (SRB) measure (defined as a measure that is absolutely continuous along the unstable manifolds) exists.

Regarding the relation of the metric entropy with the Lyapunov exponents, Ruelle proved (20) that :

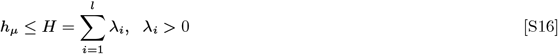

The equality, known as the Pesin identity, was shown to hold true for SRB measures. If the system is hyperbolic, the tangent space to the trajectory can be decomposed in the direct sum of its linear stable and unstable subspaces ^†^. Hyperbolicity implies the existence of an SRB measure, and that the Pesin identity and the Kaplan York conjecture hold. Nevertheless, ruling out the hyperbolicity of the system allows for no statement regarding the existence of an SRB measure. Keeping in mind that the measures derived from the Lyapunov spectrum are upper bounds and that no rigorous statement can be made in favor of the equalities, we will nevertheless refer to them as the metric entropy and the attractor dimension.

#### B. Lyapunov spectrum of delayed spiking neuronal networks

Given a neuronal model *F* (*V*_*i*_) and a model for the SCA that are exactly solvable, the network can be propagated between spikes and updated by Eqs. S10 and S7. From these equations, a Jacobian L(***x***(*n*)) at each spike time can be obtained, from which the estimation of the Lyapunov dimension *D* (Eq.S15) and the metric entropy *H* (Eq.S16) are possible via QR decomposition.

##### B.1. Derivation of the single spike Jacobian

In the following, we will focus on the derivation of 𝕃 (***x***(*n*)) for the delayed system of neuronal types that allow for a phase representation. In this case, the Jacobian elements can be written in terms of the phase response curve (PRC) (defined in Eq. S10 for the QIF case).

When a SCA with associated variable *ξ*_*m*_*∗*, spikes at *τ*_*s*+1_ ∈ [*t*_*s*_, *t*_*s*+1_], the soma units that are not postsynaptic to the axon will evolve independently of when the spike is exactly, following 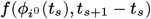 and the ones that are postsynaptic to it will be both propagated and updated 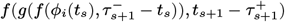 (see Eq. S10).

Equivalently, when a spike is emitted by a neuron, it will be received only by its own associated SCA. The effect of the incoming spike on the dynamics of the SCA will then be 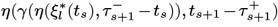. SCA that are not post-synaptic to the spiking neuron will evolve with 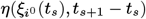 (see *Methods*).

In order to calculate the single spike Jacobian, we first summarize the equations for the phase neuron and the SCA. We define:

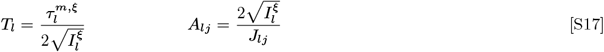

Where the supra-index *ξ* was added to emphasize that those constants are only meaningful for SCA.

The propagation and update functions for the neurons *f* and *g*, and for the SCA, *η* and *γ*, for a network of QIF neurons with QIF SCA’s are:

###### Soma equations

Propagator

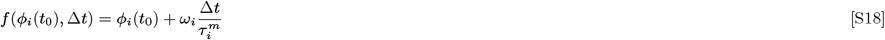

With derivatives

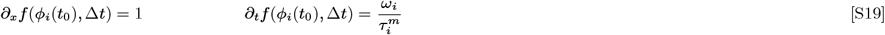

Update function

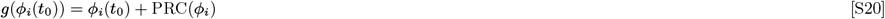

With derivatives

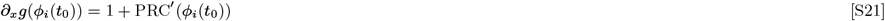

Where the phase response curve was defined in Eq. S10.

###### SCA equations

Propagator

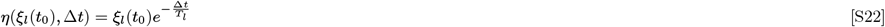

With derivatives

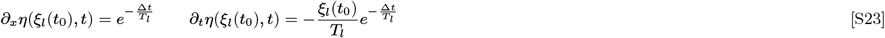

Update function

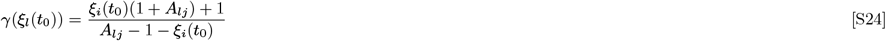

With derivatives

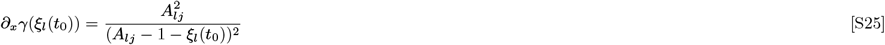

For a generic neuron model with state variable *x*, that evolves with function *f* and updates with function *g*, the derivative with respect to some other neuron variable (possibly defined by a different neuron model) *y* can be written as:

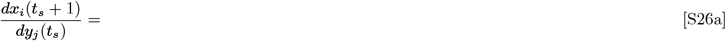

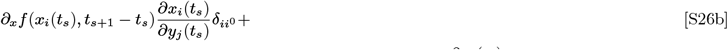

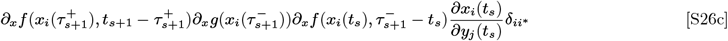

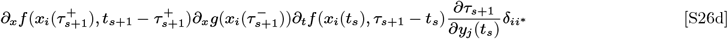

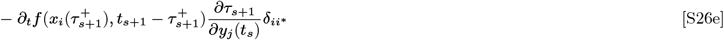

Where *i*^0^ is the index corresponding to the non-postsynaptic neurons, *i*^*∗*^ for the postsynaptic ones and the spiking neuron *j*^*∗*^. Note that if the spiking neuron is reset, then only the last term (Eq. S26e) survives given that 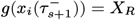 and therefore has null derivative. The term 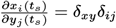 and 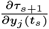 can be extracted from the fact that *f* (*y*_*j*_*∗* (*t*_*s*_), *τ*_*s*+1_ − *t*_*s*_)) = *X*_*T*_ and therefore:

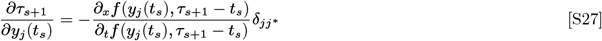

In the case we are analyzing here,

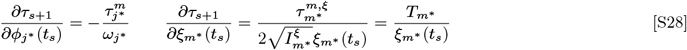

The terms of the Jacobian for the delayed system can be summarized as follows:

###### Non-postsynaptic somas 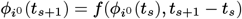

Contribute to the Jacobian only with diagonal terms from Eq. S26b

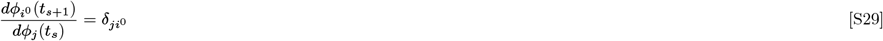

###### Postsynaptic somas 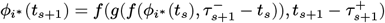

Contribute with non diagonal terms (when receiving a spike from a delayer SCA *ξ*_*m*_*∗*, from Eq. S26d and S26e

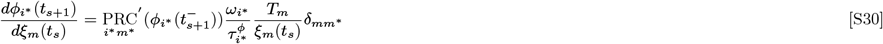

###### Spiking neuron 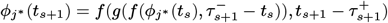

Only one term in the Jacobian, from Eq. S26e, Eq. S26c and S26d vanish

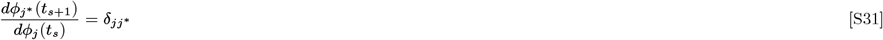

###### Non-postsynaptic SCA 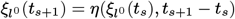

Contribute with diagonal terms from Eq. S26b

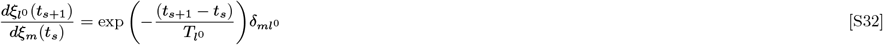

###### Postsynaptic SCA 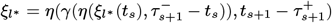

Contribute with non-diagonal terms (when receiving a spike from a neuron *ϕ*_*j*_*∗* ) from Eq. S26d and S26e

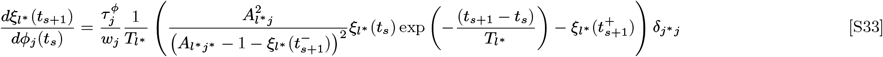

###### Spiking SCA 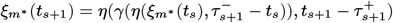

Only one term in the Jacobian, from Eq. S26e. Eq. S26c and S26d cancel one each other

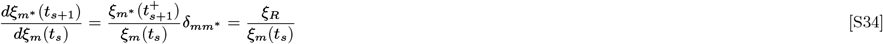

If one chooses *ξ*_*R*_ = 0, then the delay introduced in the network is exact, and the equation vanishes 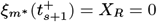 for the *delayer* SCA. In this case, the Jacobian is going to be singular. If the reset value is small but nonzero, an invertible Jacobian is obtained. In this last case, is necessary to ensure that the reset value is passed by the singularity at *ξ*_*l*_ = 1 − *A*_*lj*_ (see S24), so an incoming spike takes finite time to drive the SCA to threshold (i.e. there is a solution for Eq. S35, see below). Then it is necessary that *ξ*_*R*_ *>* 1 − *A*_*lj*_.

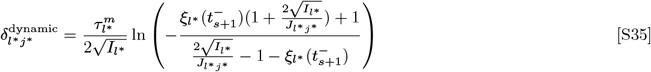

#### C. The direct method for the estimation of the first LE

We computed the first Lyapunov exponent via direct method in order to numerically corroborate the equations derived in the previous subsection. For this, after a long warm-up, we perturbed the network by adding a random vector of norm *ϵ*. After a short simulation of *T* = 100 ms the norm *β* of the difference between the perturbed and the unperturbed final states, simulated separately, is stored. The iteration of this procedure N times leads to the estimation of the first Lyapunov exponent via the following formula:

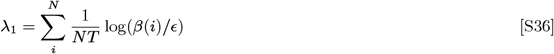

The comparison between the direct method and the one derived in the above subsection are shown in Figure S8**b**, for *ϵ* = 10^−10^ and N=5000, showing good agreement.

### 3. Delayed rate networks

We study two examples of rate networks. The activity *x*_*i*_ of each unit *i* is defined by

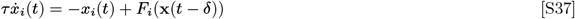

where *τ* is the time constant, *F*_*i*_ is a scalar continuous function, **x**(*t*) = *{x*_1_(*t*), …, *x*_*N*_ (*t*)*}* is the vector of all activities and *δ* is the interaction delay.

#### A. Classic rate network without Dale’s law

We modify a classic network of *N* rate units originally studied by (21) to incorporate delays. In this case,

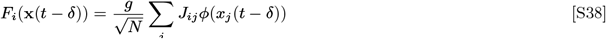

where *ϕ* is a nonlinear input-output function we choose to be *ϕ*(*x*) = tanh(*x*), *g* is a gain parameter, N is the number of neurons and *J*_*ij*_ is the connectivity matrix whose elements are Gaussian distributed with zero mean and unit variance. In the case of *δ* = 0, the system has a stable trivial fixed point for *g <* 1. At *g* = 1, the system looses linear stability and a chaotic attractor is created. Because the fixed point is the same for each neuron, the linearized delayed system can be studied by proposing a solution of the form 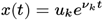, where *u*_*k*_ are the eigenvectors of *J*. The relationship between the eigenvalues of J, *µ*_*k*_, and the decay rates *ν*_*k*_ determining the linear stability of the delayed system is given by (see also (22–24)):

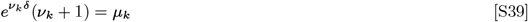

Then, for each mode, the stability is given by the solutions of *H*(*ν*) = *ν* + 1 − *µe*^−*νδ*^ = 0. The stability boundary is no longer a line at *g* = 1 but is defined by off-centered Archimedean spirals (23) that always contain the *g* = 1 point. If the elements of *J* are correlated, then the eigenvalue distribution of *J* will be ellipsoidal (25) and system can loose stability via intersections of the stability spiral with the eigenvalue spectrum giving rise to limit cycle oscillations (24). Here we focus on the chaotic regime, for *g >* 1, and for simplicity we look at uncorrelated configurations of *J* but those seem not to affect our results. We notice that the mean field theory developed in (21) would not be changed by including delayed interactions, and therefore we do not expect any change in the dynamics.

#### B. Inhibitory rate network in the balanced state

Following Ref. (26), we study a network of *N* inhibitory units which we modify to incorporate delayed interactions. In this case,

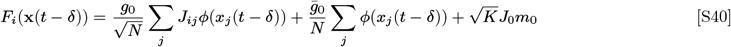

Where *m*_0_ is the target mean rate, and *J*_*ij*_ is defined as in the classic SCS case. The nonlinearity in this case is given by a threshold linear function *ϕ*(*x*) = [*x*]_+_. The size of the network N, the sparsity K and the strength of the inhibitory connections *J*_0_ are related to the variables *g*_0_ and 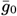 by 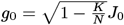 and 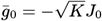.

#### C. Small delay approximation

We study the system above by making a small delay approximation. We define 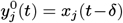. Then, 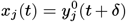 can be approximated by a Taylor expansion of order 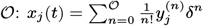. Each time derivative of 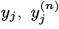, defines a new variable we call 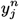. The infinite dimensional system defined in Eqs. (S38, S40) can then be approximated by the 𝒪 + 1 dimensional system given by

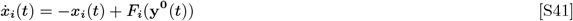

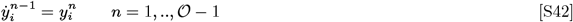

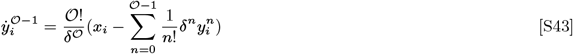

Where 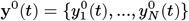.

We notice that the small delay approximation, is nothing more than expanding the term 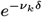 in Eq. S39 in powers of the delay *δ*. When comparing the eigenvalues of the exact delayed system against with the eigenvalues of the system under the small delay approximation, we therefore compare the solutions 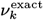 of Eq. S39 with the solutions 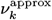 of

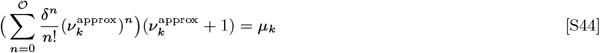

The difference among these two is shown in Figure S13**a**.

### 4. Ergodic theory of chaos for weakly delayed rate networks

Defining the N dimensional matrix 𝔸 with components 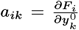, at each point in time, the Jacobian 𝕃 of the (𝒪 + 1)N system can be written as:

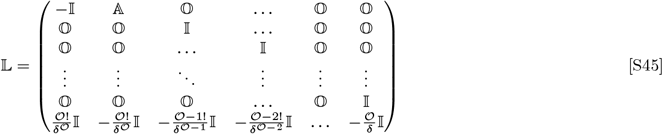

where 𝕀 is the N dimensional identity matrix. Concretely, for the delayed SCS system defined in Eq. S38 we have

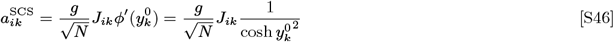

whereas for the purely inhibitory network

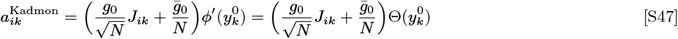

Defining now the vector **z**(*t*) = **x**(*t*), **y**^0^(*t*), **y**^1^(*t*), …, **y**^𝒪−1^(*t*), the evolution of the system defined in S41, can be re-written by defining G such that *ż*_*i*_ = *G*_*i*_(**z**).

The calculation of the Lyapunov exponents can be done following Engelken et al. (27).

**Fig. S1.**
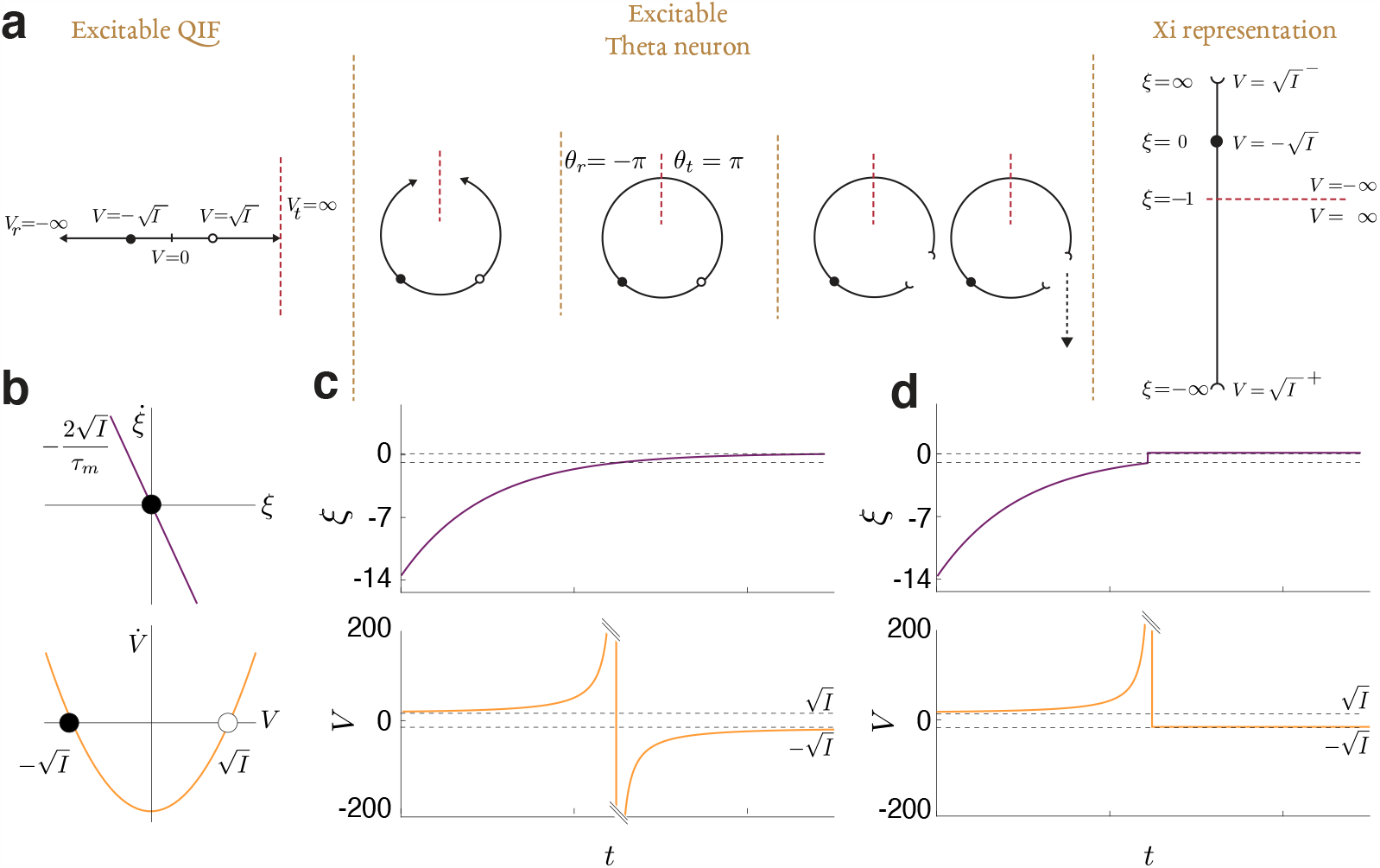
Single compartment axonunit. **(a)** Three possible representation for the dynamics of the QIF neuron. (Left) A quadratic integrate and fire (QIF), has two fixed points at 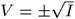, an stable one (black) and an unstable one (white). This neuron is solvable analytically either in the tonic spiking regime or the excitable regime but nor in both. The neuron model diverges in finite time, emitting a spike. (Middle) A classic variable transformation was introduced by (28) and is known as the theta neuron. (Right) We introduce a variable transformation, the *ξ* representation, tailored to the SCA. This representation is analytically solvable. The SCA at rest in *ξ* = 0 is pulled towards −∞ when receiving a spike, and evolves towards zero, emitting a spike at *ξ* = −1 **(b)** Phase space of the SCA *ξ* (Top) and the canonical QIF representations (Bottom) **(c)** Time evolution without reset to the fixed point *ξ*_*r*_ = *ξ*_*t*_ = −1. When the *ξ* variable crosses -1 (Top), in the voltage representation the voltage diverges (Bottom) **(c)** Time evolution resetting the SCA to its stable fixed point i.e. *ξ*_*r*_ = 0 in both representations.

**Fig. S2.**
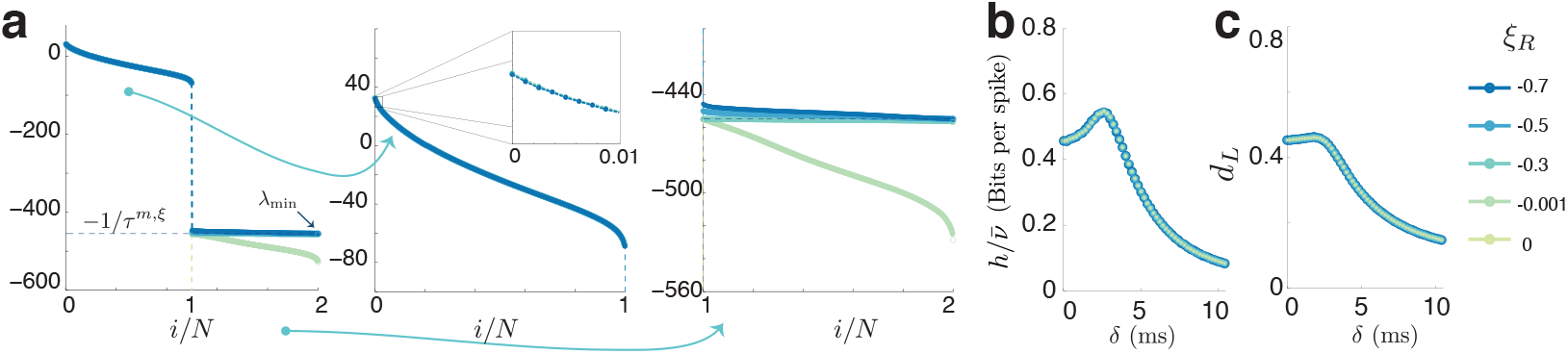
Dependence on the reset values of the single compartment axon. (see Eq. S35 and *Methods*); *ξ*_*r*_ = 0 corresponding to the exact delay framework.**(a)** Full spectrum for different values of the reset. The spectrum is composed of two branches (left). The first one (middle)characterizes the biologically relevant degrees of freedom (up to *i* = *N*, where N is the number of soma units in the network). The second branch (right) is defined by the time it takes for the axon to relax back to its fixed point. In the “exact delay” framework where *ξ*_*r*_ = 0, that time constant is zero and therefore the second branch of the Lyapunov exponents diverges to−∞. If the reset point is close to the fixed point the SCA will relax faster than what its dictated by its time constant *τ* ^*m,ξ*^. As can be seen, the first part of the spectrum seems to be independent on the choice of *ξ*_*r*_, given that the time constant is chosen such that they dynamics are not visibly affected. This results on identical dependencies of the normalized metric entropy *h* **(b)** and attractor dimension **(c)** of the value of the delay.

**Fig. S3.**
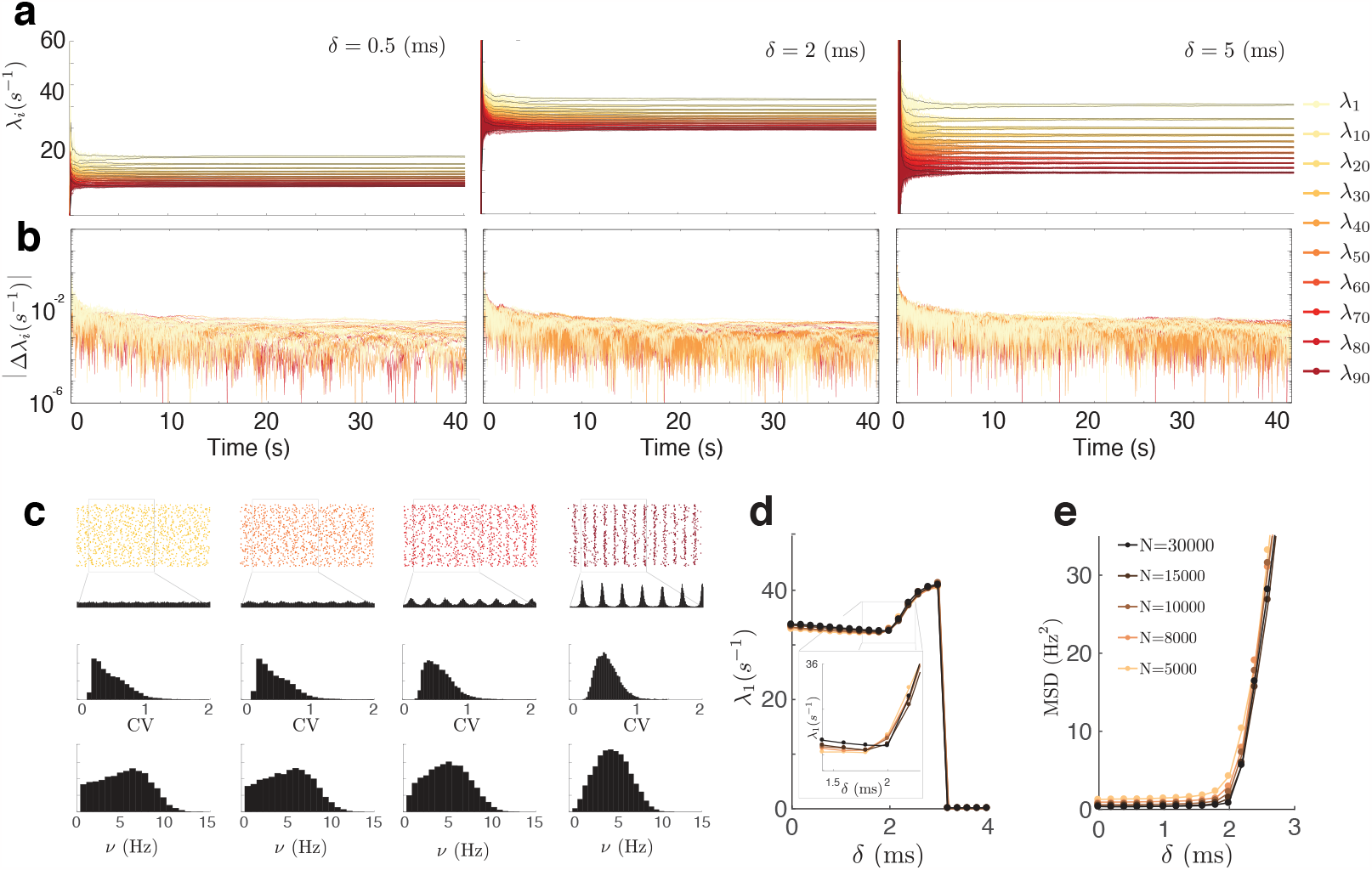
Convergence of the Lyapunov spectrum and finite size effects in the first Lyapunov exponent. **(a)** Example traces of Lyapunov exponents for different values of the delay. **(b)** Difference between different different LE traces for different instantiations of the random connectivity |Δ*λ*_*i*_| get smaller with time. Over the course of seconds simulations the LE converge to its true value. **(c)** Distributions of coefficient of variations and firing rate distributions for the example rasters in Figure 3 of the main text. **(d)** First Lyapunov exponent as a function of the delay for different network sizes N. The larger the network the clearer the downward dependence of the first exponent with the delay before the transition. **(e)** MSD indicating that the increase in the exponent arises from the mean field Hopf bifurcation.

**Fig. S4.**
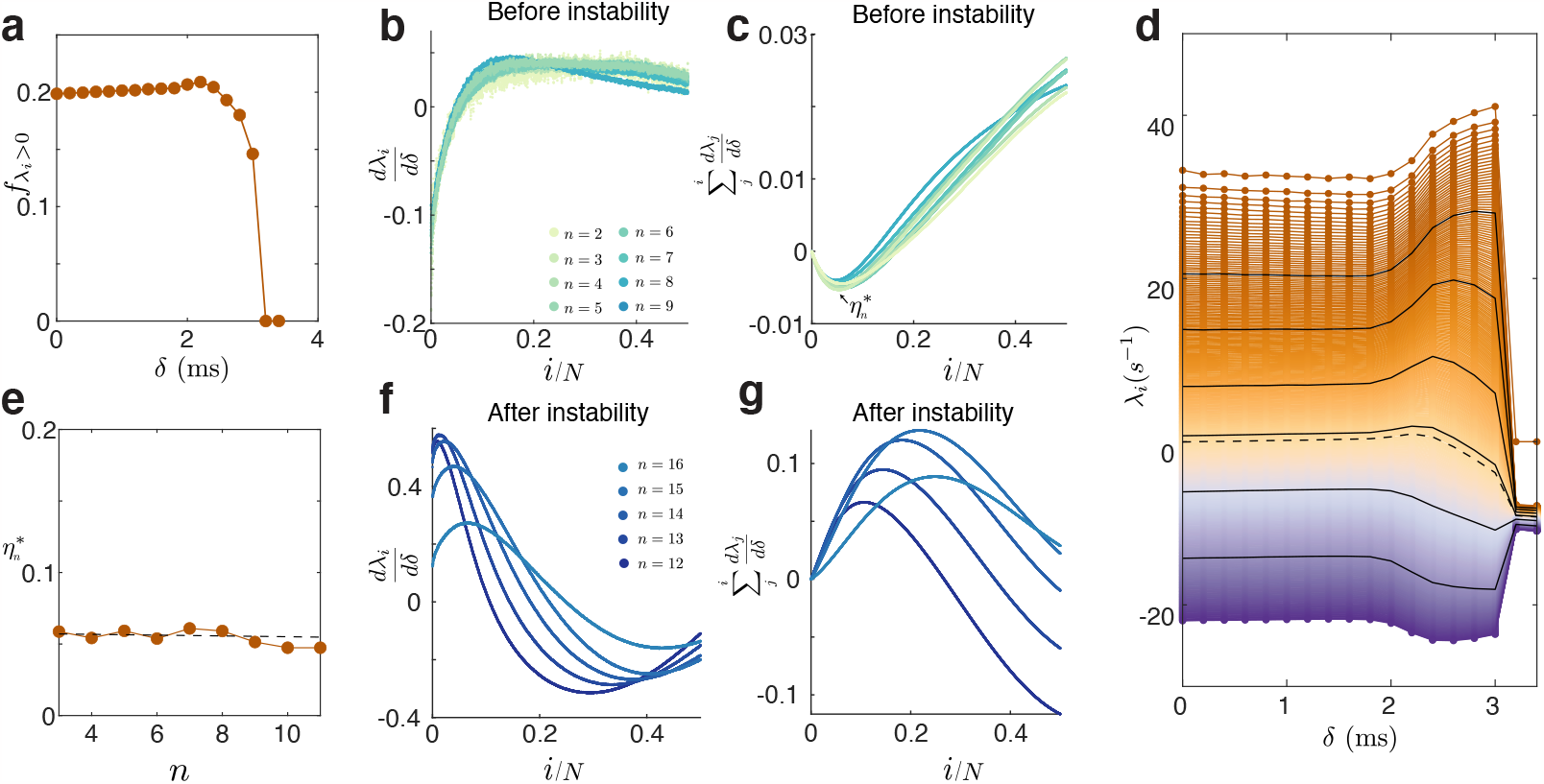
Structure of the Lyapunov Spectrum (K=50). **(a)** Fraction of positive Lyapunov Exponents. **(b)** Slope of the *λ*_*i*_ computed by 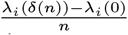, where *n <* 10, where the transition occurs and *λ*_*i*_(0) is the spectrum of the non=-delayed system **(c)** Cumulative sum of the magnitude computed in (b). The minimum 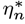 indicates the fraction of LE that have an initially negative dependence with the delay as computed by taking the *n* th difference. Approximately the first 5% LE decrease their magnitude with increasing delay before the transition. **(d)** All biologically-relevant Lyapunov exponents as a function of the delay for a single realization of the random connectivity. After the transition at *δ*_*c*_ = 3 ms the maximum lyapunov exponent vanishes (the trivial exponent is present because of the translation invariance of the system) and the rest of the spectrum relaxes to the inverse of the membrane time constant *tau*_*m*_ = 10 ms. After the instability, all positive Lyapunov exponents increase in magnitude (the LE that was initially zero is in dash). Negative LE decrease in magnitude. **(e)** Fraction of LE that have a negative slope with the delay before the transition as a function of the *n*th difference. **(f)** Slope of the *λ*_*i*_ computed by 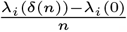 for n after the oscillatory transition. Right after the transition most LE have a larger value that in the non-delayed case **(g)** Cumulative sum of the magnitude computed in (f)

**Fig. S5.**
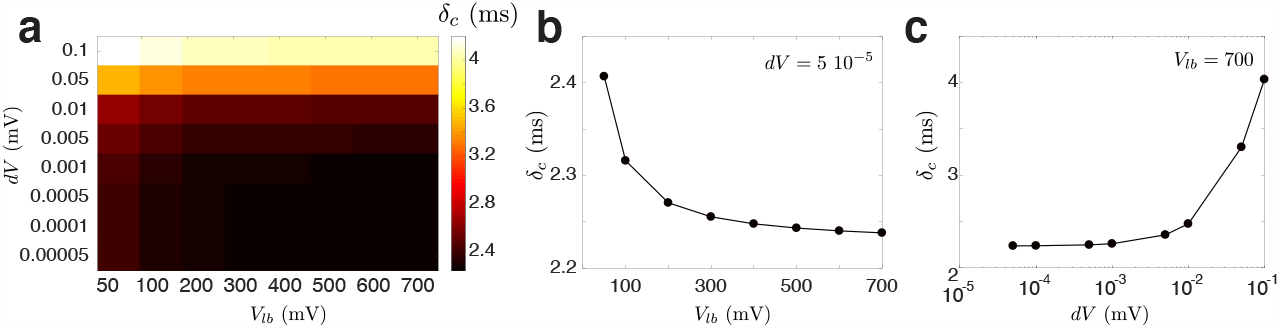
Numerical precision of the Richardson’s method. **(a)** The threshold-integration method developed by Richardson (11) involves integrating the first derivative of the voltage probability distribution and that of the current backwards from threshold to a lower bound of the voltage. The QIF neuron used here does not have a threshold or a reset value, given that it diverges in finite time and therefore the critical delay computed with this method will depend on the choice of the integration bounds. Empirically nevertheless we see that the estimation changes very little after a certain threshold value. Critical delay for a network of K=50, target firing rate =5 Hz, and other parameters as default, as a function of *dV* (the integration step) and the value of the threshold chosen for integration. **b-c)** Cross sections of a)

**Fig. S6.**
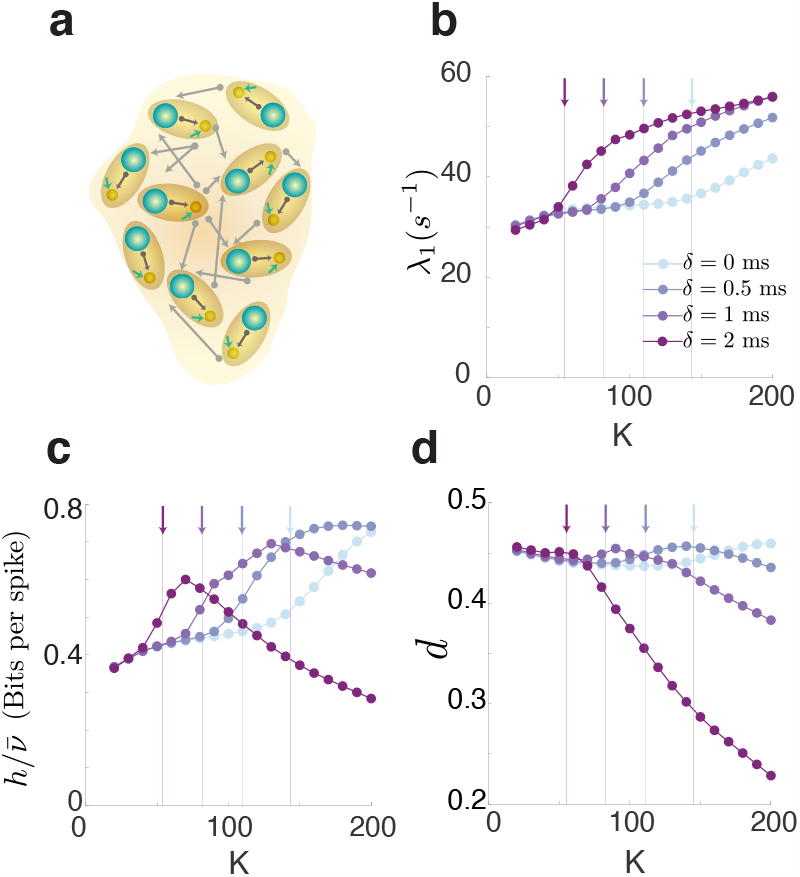
Transition to collective oscillations as a function of the average indegree K. Because of the balanced-state scaling in which the connection weights scale as 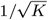, increasing K also makes each connection weaker. **(a)** Inhibitory network with delays. **(b)** Maximum Lyapunov exponent as a function of K for different values of the delay. The critical value of the indegree at which the asynchronous irregular state loses stability to a collective oscillation is computed as in Figure 4 of the main text, and indicated by the vertical arrows in matching colors.**(c)** metric entropy as a function of K for different values of the delay. **(d)** Attractor dimension as a function of the delay. Even if the attractor dimension decreases with K up to the oscillatory instability, it suddenly increases at the transition. All measures of chaos increase at the oscillatory transition.

**Fig. S7.**
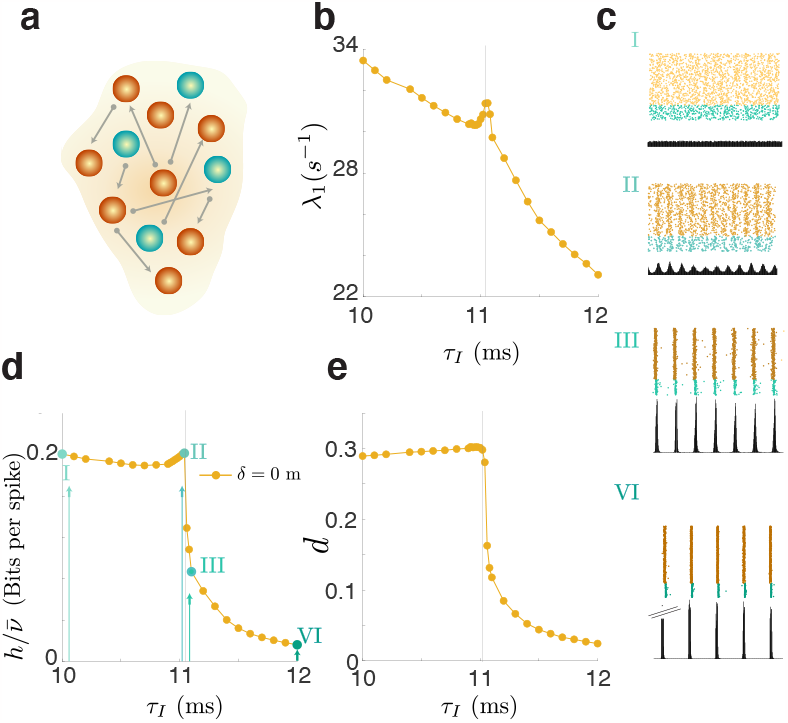
**(a)** Non delayed E-I network in the balanced state. The asynchronous irregular state loses stability with increasing inhibitory time constant as described theoretically in (2). We see that at the onset of collective oscillations, all chaotic measures increase. **(b)** Maximum Lyapunov exponent as a function of of the inhibitory time constant *τ*_*I*_ **(c)**. Raster plots for different values of the inhibitory time constant. **(d)** metric entropy as a function of *τ*_*I*_ **(e)** Attractor dimension as a function of *τ*_*I*_.

**Fig. S8.**
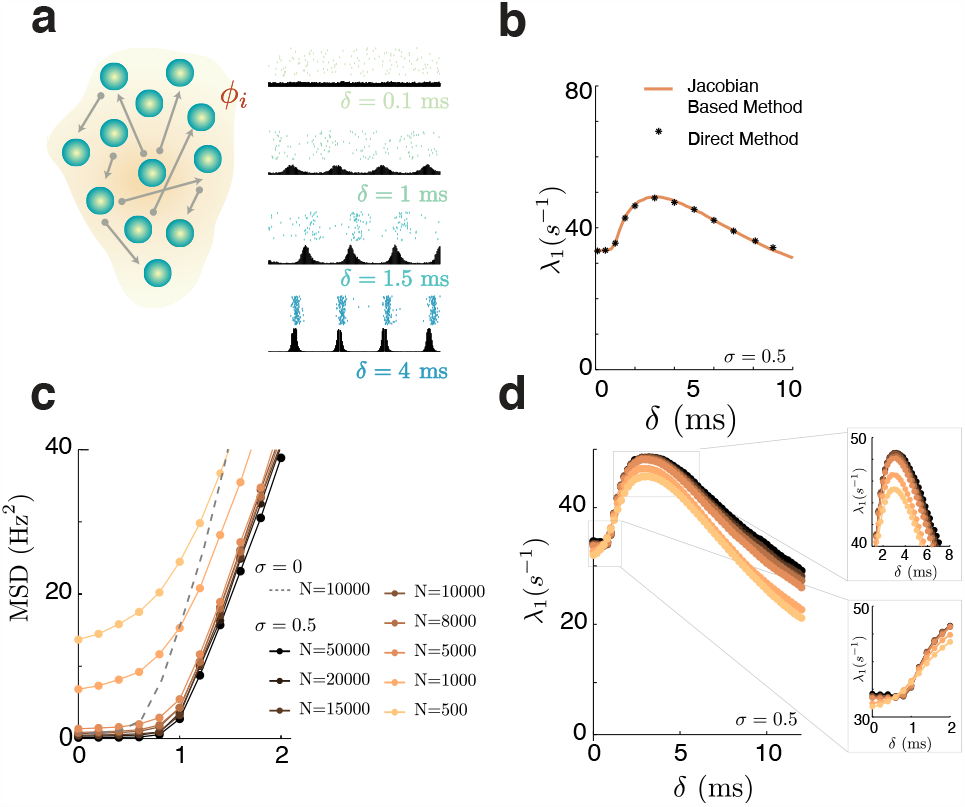
Single cell heterogeneity perpetuates chaos. **(a)** Network of inhibitory neurons with heterogeneity and raster plot of the network activity for different delays, observe the difference in the spiking pattern after the first oscillatory transition (middle bottom panel) and after the second one (bottom panel). **(b)** Direct computation of the maximum LE, by perturbing the trajectory numerically is compared with the method developed here (*N* = 5000) showing good agreement. **(c)** MSD and **(d)** Maximum LE for different network sizes as a function of the delay for an example heterogeneous network of *σ* = 0.5. The dashed line in **(c)** indicates the MSD for an homogeneous network for reference. The maximum exponent converges to a size independent value which increases with the network size, indicating that finite size network fluctuations only lead to an under-estimation of the chaotic nature of the circuit.

**Fig. S9.**
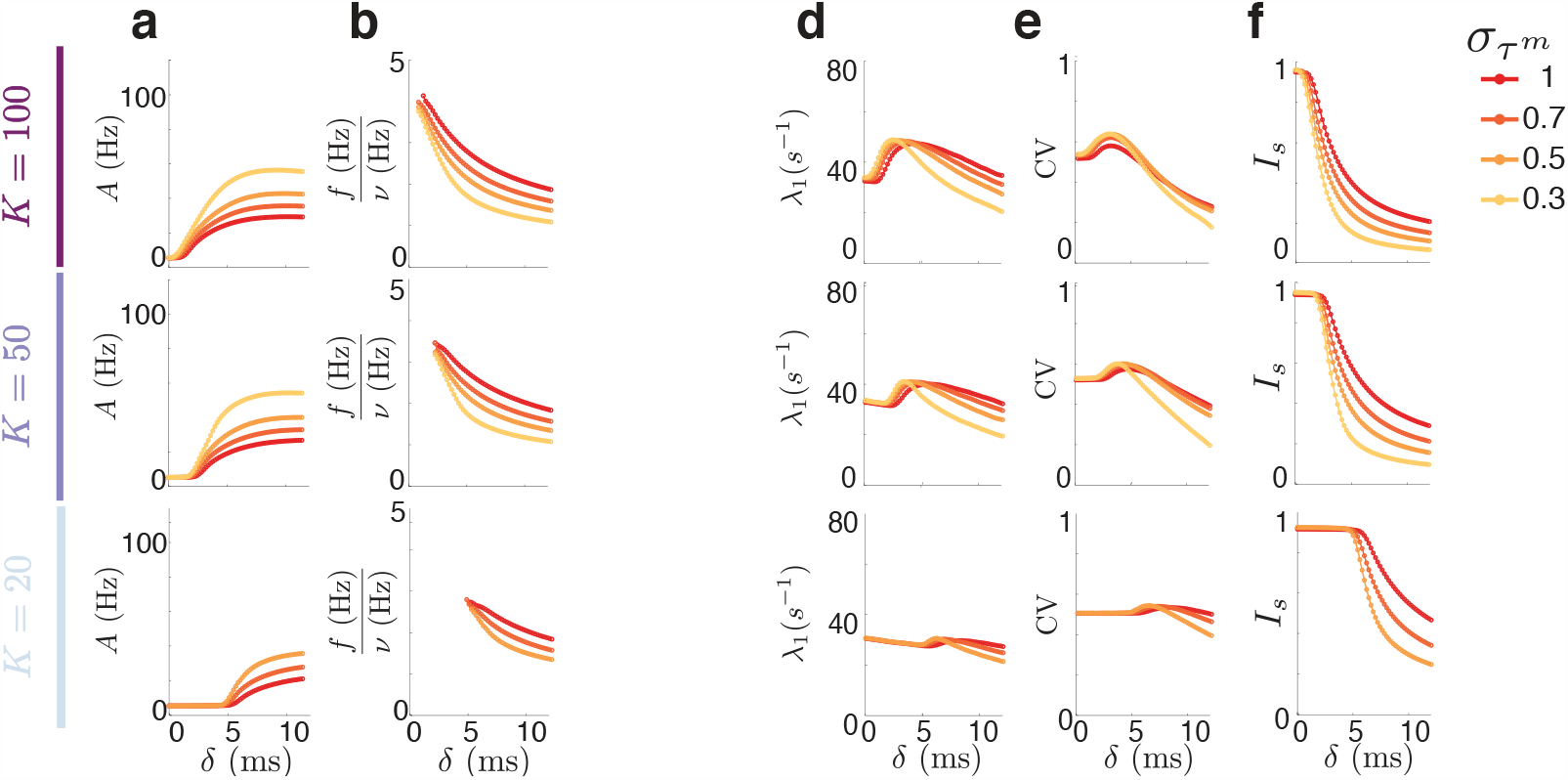
Impact of heterogeneity. **(a)** Amplitude and **(b)** frequency of the population oscillation as a function of the delay for different values of membrane time constant heterogeneity (uniform distribution of mean 10ms and standard deviation 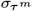). **(d)** The maximum Lyapunov exponent and the **(e)** coefficient of variation have similar dependencies to the delay. **(f)** 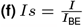 is the ratio between the input current to the neurons needed to reach the target firing rate compared to what its predicted theoretically from the balanced state condition Eq. S3

**Fig. S10.**
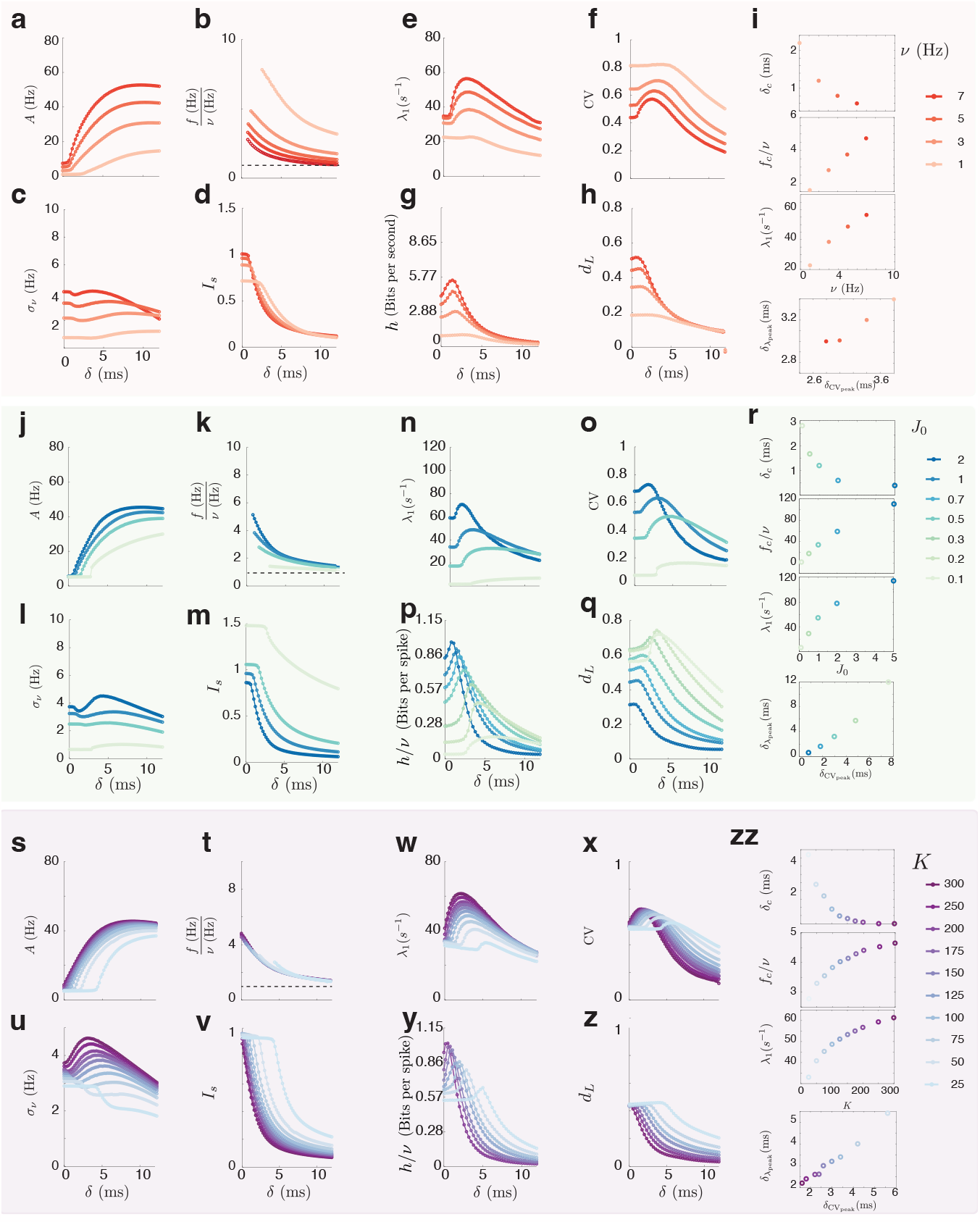
Robustness of the findings. **(a-i)** Robustness for different values of the mean firing rate **(a)** Amplitude of the time dependent population rate (also multi-unit activity, MUA) as a function of the synaptic delay *δ*. **(b)** Frequency of the oscillation vs *δ*. The values depicted are shown only when the MUA amplitude departed from baseline **(c)** Standard deviation of the firing rate distribution vs *δ*. (d) 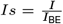 is the ratio between the input current to the neurons needed to reach the target firing rate compared to what its predicted theoretically from the balanced state condition Eq. S3 This number quantifies a departure from the balanced state equation. **(e)** Maximum Lyapunov Exponent vs *δ*. (f) Coefficient of variation vs *δ*. (g) metric entropy vs *δ*. Notice that its not normalized by the firing rate and the units are then bits per second. (h) Lyapunov Dimension. Parameters: Default parameters as in table 1 (see *Methods*), except for panels (g-h) where N=2000 and the changes indicated in the figure. **i)** Top: Dependence of the critical delay with the mean rate *ν*. Top-Middle: Frequency at the onset of collective oscillations as a function of *ν*. Middle Bottom: Maximum value of the maximum Lyapunov exponent as a function of *ν*. Bottom: value of the delay at which the maximum lyapunov exponent peaks vs the value of the delay at which the coefficient of variation peaks. **(j-r)** Robustness for different values of the connectivity strength *J*_0_. Panels identical to (a-i). **(s-zz)** Robustness for different values of K. Panels identical to (a-i).

**Fig. S11.**
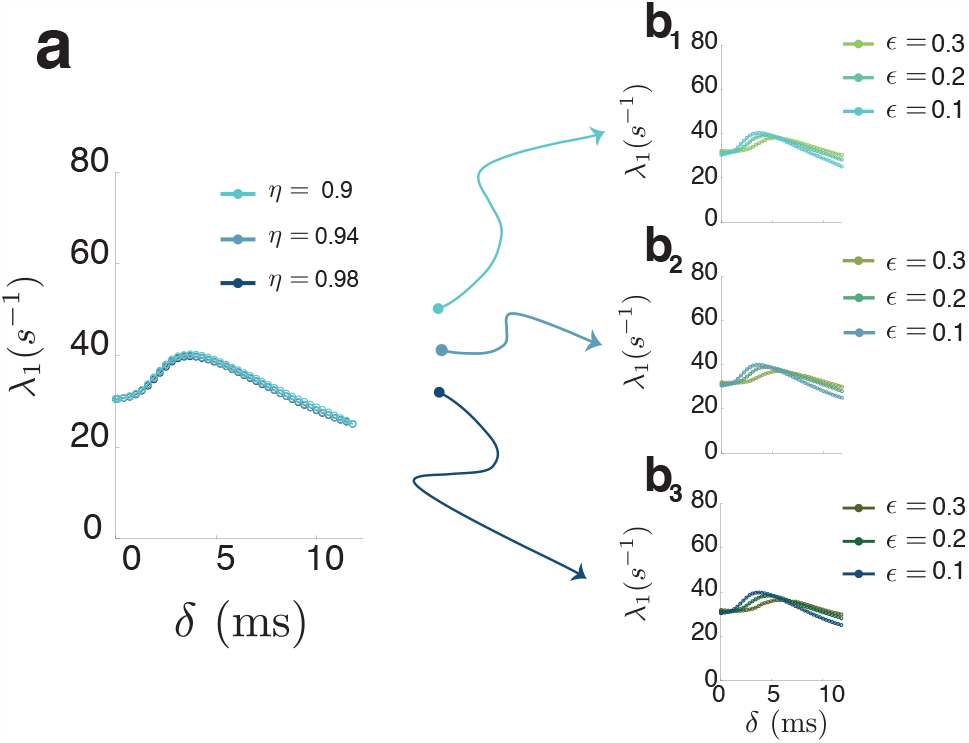
Dependence of the maximum Lyapunov exponent on the intensity of the EI loop. **(a)** The maximum Lyapunov exponent is invariant under small changes in the excitation ratio parameter *η* (see Eq. A). **(b)** Different values of the feedback loop *ϵ* delay the transition to collective rhythms and push the peak of the exponent towards larger delays. Parameters: Default parameters except for N=5000

**Fig. S12.**
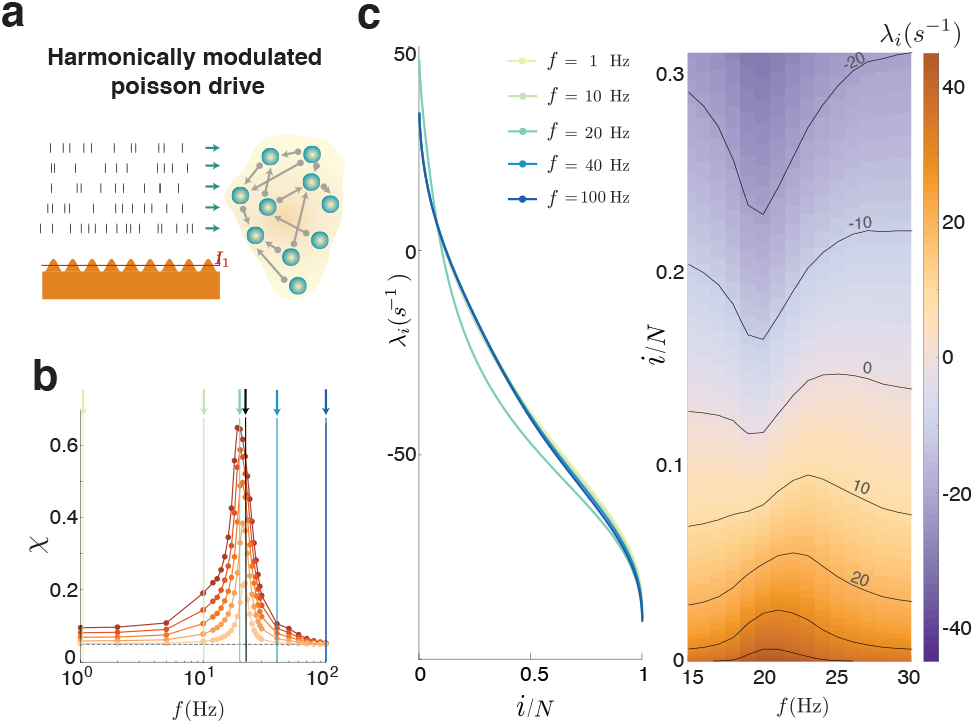
**(a)** Harmonically driven case **(b)** Synchronization index as a function of the frequency of the harmonic drive *f* **(c)** Entire spectrum of the LE of the driven network for different values of the frequency of the harmonic input drive (left), and a third of the spectrum for frequency of the drive *f* between 15 and 30 Hz. Notice how when the maximum Lyapunov exponent starts to increase, the negative exponents become more negative, resulting unexpectedly in a decrease instead of an increase of the attractor dimension for frequencies smaller than the critical one.

**Fig. S13.**
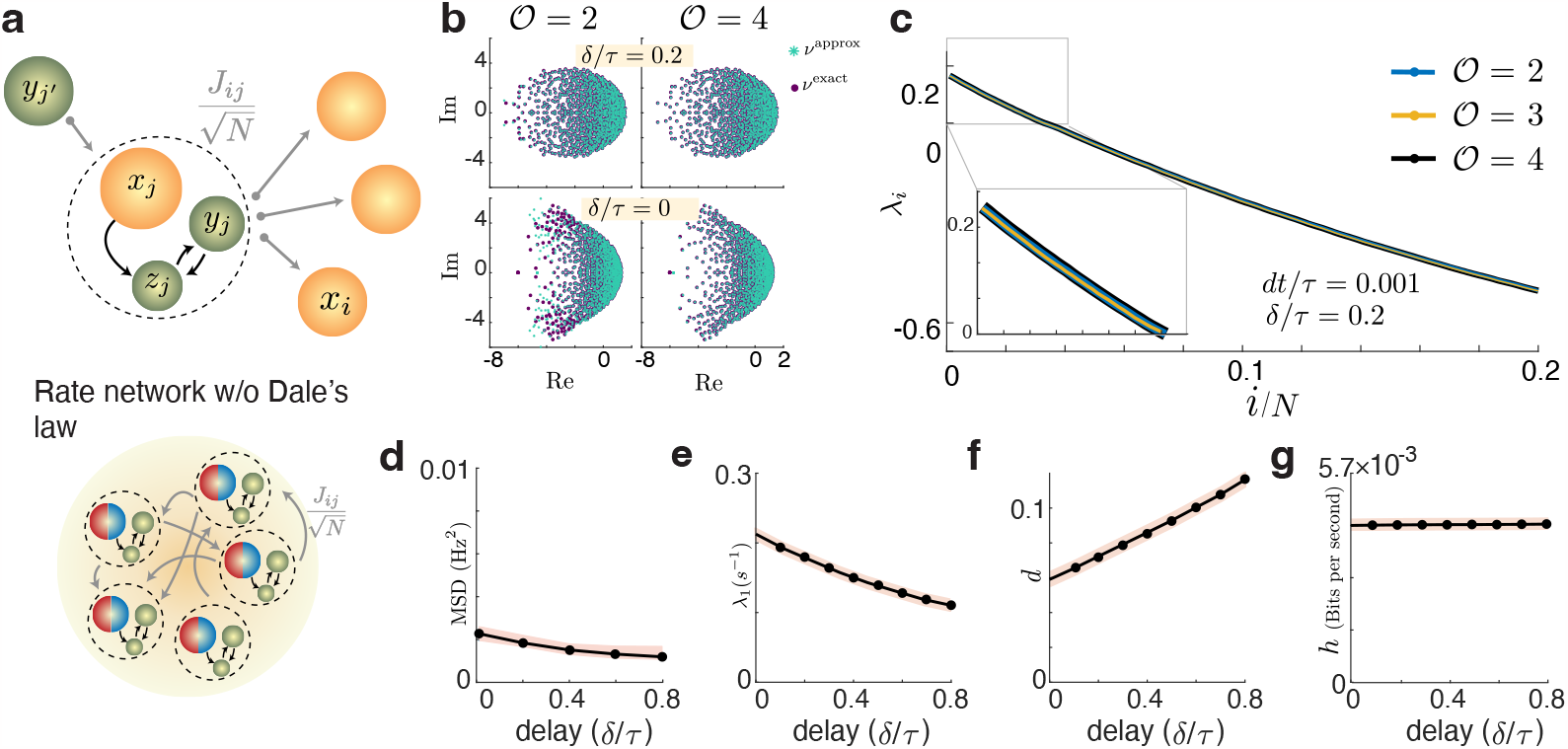
Classic rate networks behave like classic scalar delayed systems. **(a)** Diagram of the small-delay approximation network. There will be 𝒪 *N* extra units for an approximation of order 𝒪 and a network of *N* neurons (see *Methods*) **(b)** Distribution of perturbation decay rates *ν*_*k*_, associated to the eigenvalue *λ*_*k*_ of the random matrix *J*, for the exact delayed system (purple) and for the small delayed approximation (turquoise) of order 𝒪 = 2 (left column) or 𝒪 = 4 (right column) **(c)** Lyapunov spectrum of the approximate system for orders 2, 3 and 4 for an intermediate delay **(d)** Mean square deviation **(e)** Maximum Lyapunov exponent **(f)** Attractor dimension and **(g)** Metric entropy as a function of the synaptic delay. Notice that the metric entropy is in bits per second.

**Fig. S14.**
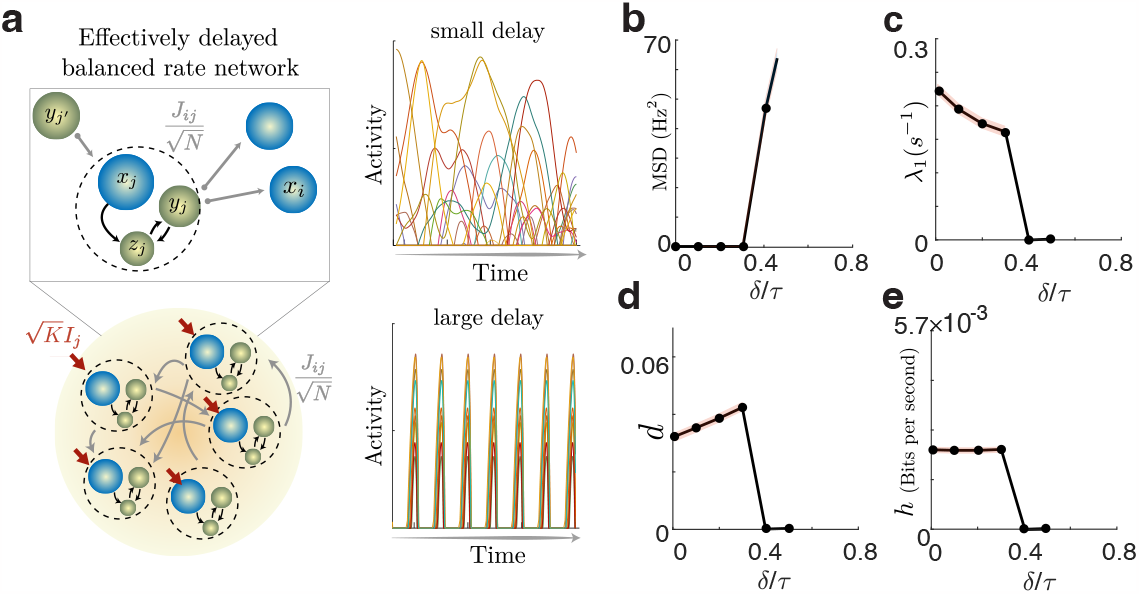
Inhibitory delayed rate networks behave like classic scalar delayed systems. **(a)** Diagram of the small-delay approximation network. There will be 𝒪 *N* extra units for an approximation of order 𝒪 and a network of *N* neurons (see *Methods*) Both the delayed rate network and the approximated “weakly delayed” networks used here have a transition from fully developed chaos (top) to clock-like synchrony. **(b)** Mean square deviation from the mean is vanishingly small before the critical delay indicating asynchronous dynamics. **(c)** First Lyapunov exponent as a function of the delay **(d)** Attractor dimension **(e)** Metric entropy.

## Footnotes

† This means that the tangent spaces are never co-linear, there is a finite angle between them, and the tangent space can be written as *TM* = *E*^*u*^ ⊕ *E*^*s*^

